# Vaginal metatranscriptome meta-analysis reveals functional BV subgroups and novel colonisation strategies

**DOI:** 10.1101/2024.04.24.590967

**Authors:** Scott J. Dos Santos, Clara Copeland, Jean M. Macklaim, Gregor Reid, Gregory B. Gloor

**Author notes:** Contributing authors.

## Abstract

The application of ‘-omics’ technologies to study bacterial vaginosis (BV) has uncovered vast differences in composition and scale between the vaginal microbiomes of healthy and BV patients. Compared to amplicon sequencing and shotgun metagenomic approaches focusing on a single or few species, investigating the transcriptome of the vaginal microbiome at a system-wide level can provide insight into the functions which are actively expressed and differential between states of health and disease. We conducted a meta-analysis of vaginal metatranscriptomes from three studies, split into exploratory (*n* = 44) and validation (*n* = 297) datasets, accounting for the compositional nature of sequencing data and differences in scale between healthy and BV microbiomes. Conducting differential abundance analyses on the exploratory dataset, we identified a multitude of strategies employed by microbes associated with states of health and BV to evade host cationic antimicrobial peptides (CAMPs); putative mechanisms used by BV-associated species to resist and counteract the low vaginal pH; and potential approaches to disrupt vaginal epithelial integrity so as to establish sites for adherence and biofilm formation. Moreover, we identified several distinct functional subgroups within the BV population, distinguished by genes involved in motility, chemotaxis, biofilm formation and co-factor biosynthesis. After defining molecular states of health and BV in the validation dataset using KEGG orthology terms rather than community state types, differential abundance analysis confirmed earlier observations regarding CAMP resistance and compromising epithelial barrier integrity in healthy and BV microbiomes, and also supported the existence of motile vs. non-motile subgroups in the BV population. Our findings highlight a need to focus on functional rather than taxonomic differences when considering the role of microbiomes in disease and identify pathways for further research as potential BV treatment targets.

## 1 Introduction

Bacterial vaginosis (BV) is a polymicrobial condition of poorly-understood aetiology and is the most common cause of vaginal concern worldwide [1]. Prevalence rates vary by country and are typically between 20-30 % [2], though reported rates exceed 50 % in some rural populations of developing nations [3]. Symptoms of the condition include a thin and potentially malodorous discharge, vaginal itching or irritation, pain on urination and/or sexual intercourse, and elevated vaginal pH; however many cases can be asymptomatic. Untreated, BV can have profound consequences, including higher acquisition rates of sexually-transmitted infections [4, 5] (including HIV and oncogenic HPV), in addition to a marked negative impact on patients’ self-esteem, feelings of shame or anxiety, reduced intimacy and relationship problems [6] and an increased incidence of pre-term birth [7]. Recommended treatment with oral or intravaginal metronidazole or clindamycin often achieves symptom resolution in the short term, but is associated with high recurrence rates of 50-80 % within a year [8].

Vaginal microbiome dysbiosis is a hallmark of BV and typically manifests as a depletion of *Lactobacilllus* spp., with a corresponding overgrowth of various obligate anaerobic species [9]. Typically, total bacterial loads in BV are one to two orders of magnitude higher than in healthy patients [10], and species from the genera *Gardnerella*, *Prevotella*, *Atopobium*, *Megasphaera*, *Sneathia*, and *Mobiluncus*– among others– predominate. Landmark studies of the vaginal microbiome using amplicon sequencing defined five canonical ‘community state types’ (CSTs), with CST IV showing a significant correlation with BV [11]. This finding has since been extensively replicated, including studies employing alternative molecular barcoding genes [12, 13] and shotgun metagenomic methods [14].

Characterising microbial gene expression in the vaginal microbiome remains an uncommon approach, largely due to the costs associated with obtaining sufficient sequencing depth for a large number of samples and the difficulty of analysis because of the very different species present in the two conditions. However, defining the vaginal metatranscriptome via RNA-seq enables a functional assessment of gene expression within the vaginal niche, in addition to taxonomic characterisation of the community. Identifying differences in actively expressed genes and pathways at the systems level between states of health and BV could further our understanding of the complex aetiology of BV or yield novel treatment targets. The initial taxonomic [11] and functional [15] explorations of the vaginal microbiome by culture-independent means represented a huge advance in BV research; however, while replication of the taxonomic differences in BV is reassuring, such studies have not yet advanced currently available treatments; although vaginal microbiome ‘transplantation’ has been explored [16]. Meta-’omics’ approaches are also limited by a lack of standardised methods, and findings can sometimes be difficult to reconcile or replicate between datasets– especially when different sample collection methods, bioinformatic pipelines, and statistical analyses have been used. Moreover, small study populations and inappropriate analyses in such studies may lead to false-positive findings which are spuriously correlated with disease states, rather than truly differential [17].

One final issue with metatranscriptomic analysis is marked asymmetry between datasets. The vaginal microbiome is known to be exceedingly asymmetric with bacterial loads differing by up to two orders of magnitude [10] in addition to a large gene content difference between conditions [18–20]. These differences cause an asymmetry in the location of ‘housekeeping’ genes during the data analysis that is not accounted for by the existing standard normalisation methods. This often results in a large proportion of all genes in the dataset to be incorrectly called as significantly different between experimental conditions [21, 22]. The issue of asymmetry can be overcome by incorporating a new statistical approach termed scale reliant inference (SRI) [17]. This new statistical approach included in the updated ALDEx2 R package [21, 23], allows researchers to build a Bayesian posterior model of the data that enables robust and reproducible inference even in the face of extreme data asymmetry.

We carried out a meta-analysis of three publicly available vaginal metatranscriptomic datasets. We identified expressed genes and functions which differentiate BV and healthy vaginal microbiomes, and which are reproducibly distinct across different study populations. Collecting and re-processing three RNA-seq datasets using a standardised workflow including annotation with the VIRGO pipeline [19], we found that *Lactobacillus* spp. and BV-associated taxa employ different strategies to evade host anti-microbial defense systems such as cationic antimicrobial peptides (CAMPs). Furthermore, we identified putative mechanisms for BV-associated species to counteract the low vaginal pH and compromise vaginal epithelial barrier integrity. We also observed at least two distinct subgroups of BV-associated metatranscriptome profiles differing by expression of motility, chemotaxis and co-factor biosynthesis genes. These findings were replicated in a large, independent dataset; various strategies for CAMP resistance and the existence of motile vs. non-motile BV sub-populations were evident among differentially expressed pathways. Overall, our findings highlight potential targets for novel therapeutic strategies in the treatment of BV, opening several avenues of investigation for future work made possible only by incorporating scale uncertainty in our analytical approach.

## 2 Results

### 2.1 Study cohort and final dataset

Five vaginal metatranscriptomic datasets were identified for meta-analysis, three of which were collected for analysis (Figure 1) and are described by [7, 24, 25]. Two other datasets exist but but were not used. The first [20] was an extended time-series survey of 39 participants and was not included as its size would not significantly increment our power. The second [26] interrogated the vaginal metatranscriptome and host response during severe SARS-CoV-2 infection; raw data for this study were not available at the accession number given in the article. Owing to this, and the potential confounding introduced by viral infection and intensive care admission, this dataset was also excluded.

**Fig 1.**
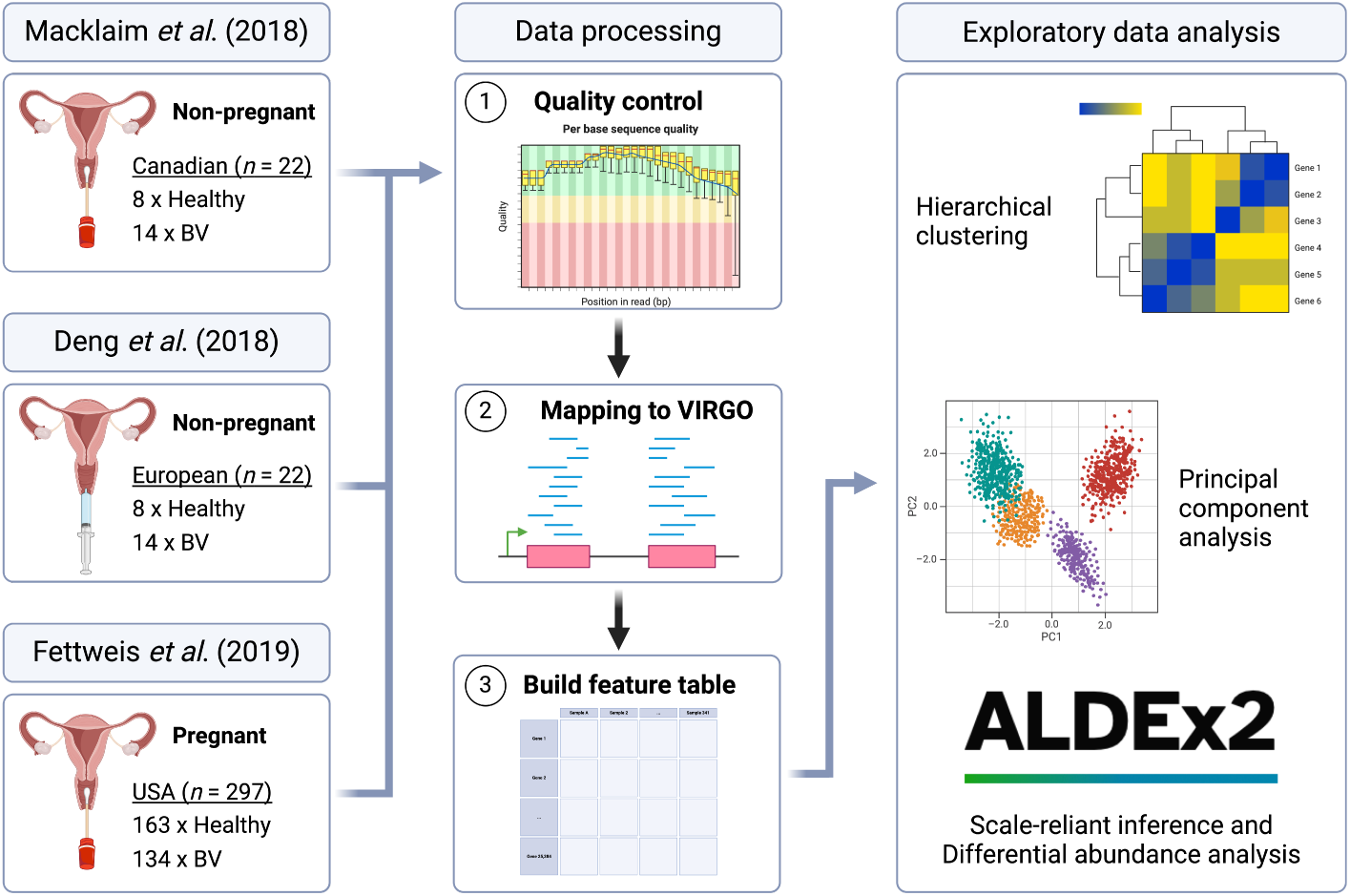
Study workflow: Raw FASTQ files were downloaded from their respective repositories and were trimmed for quality and length. Reads mapping to GRCh38 and T2T human reference genomes were discarded as presumptive human contamination; the remainder were mapped to v138.1 of the SILVA rRNA database, and any aligned reads were also discarded. The remaining reads were mapped against the VIRGO catalogue of ∼1 million non-redundant, microbial genes from the human vagina [19]. Feature tables for each dataset were constructed separately and batch-corrected [28] when used in combination. These data tables served as input for all downstream analyses using the scale reliant inference (SRI) approach to normalisation [17] and the expected values of Bayesian posterior models of the data were used as starting points for all exploratory data analysis and statistical methods.

The three study populations analysed exhibited considerable variation, suggesting that any findings reported would be general and not specific to any one dataset. Macklaim *et al*. (London dataset [24]) studied a convenience sample of 22 non-pregnant, volunteers from Ontario, Canada with the aim of identifying functional differences between states of health and BV. Deng *et al*. (Europe dataset [25]) studied 22 non-pregnant, symptomatic BV patients from Germany who underwent metronidazole treatment. These were grouped by treatment success (healthy) or failure (BV). Demographically, 84% of the women were Caucasian and the remainder had some African ancestry [27]. Fettweis *et al*. (Virginia dataset [7]) collected 297 vaginal metatranscriptome samples from pregnant volunteers as part of the MOMS-PI dataset through the integrative Human Microbiome Project in a USA-based, prospective, longitudinal cohort study aiming to ascertain the effect of the vaginal microbiome on preterm birth. Samples from London and Europe datasets were obtained primarily from Caucasian women, whilst those from the Virginia dataset were primarily from African-American women. A total of 341 metatranscriptome samples were included in the meta-analysis; median post-processing read counts were relatively similar across London and Europe datasets but higher in the Virginia dataset (Table 1).

**Table 1.** Pre-and post-processing read counts for the three datasets included in the meta-analysis. H, healthy; BV, bacterial vaginosis.

| Dataset | No. samples |  |  | Median no. reads |  |
| --- | --- | --- | --- | --- | --- |
|  | H | BV | Total | Raw | Post-processing |
| London [24] | 8 | 14 | 22 | 14,479,786 | 11,107,052 |
| Europe [25] | 8 | 14 | 22 | 16,584,536 | 8,096,724 |
| Virginia [7] | 163 | 134 | 297 | 46,368,206 | 19,967,915 |

### 2.2 Applying scale to healthy and BV metatranscriptomes

The vaginal microbiome of healthy and BV subjects differs greatly in both composition and scale; the BV microbiome is both more complex and has about two orders of magnitude more bacteria per collected volume [10]. Commonly used data normalisations assume that the majority of the parts being sampled are relatively invariant in the sampled environment. To account for this asymmetry, we used a new statistical model for high throughput sequencing datasets called scale-reliant inference (SRI) [17]. SRI allows the explicit modelling of both the amount of variation in the underlying datasets and the proper location of marker functions, and this feature is incorporated into the latest version of ALDEx2 [21, 29].

Applying SRI through ALDEx2 generates a full Bayesian posterior model of the underlying environment from the observed table of counts by incorporating uncertainty in both the observed count table and the count normalisation (i.e. scaled log-ratio transformation). This approach prevents converging on a precise but incorrect set of differentially abundant features [17]. The model is built by explicitly acknowledging two *a priori* assumptions about the table of counts. Firstly, that the expression of genes encoding standard housekeeping functions required by all organisms should be relatively invariant between groups; that is, the location of housekeeping functions should exhibit no difference between groups. We found that the naΪve estimate of scale produced by ALDEx2 had an offset of about 8-fold in scale that resulted in the location of housekeeping genes being off the location of no difference. We found that adding a 1.15-fold offset to the scale parameter was able to centre the housekeeping functions (Methods, and Supplementary Figure S1). This is biologically intuitive in that the small difference in scale needed to centre the housekeeping functions indicates that they can be assumed to be relatively invariant. We therefore applied this 15 % offset as part of all differential abundance analyses conducted with ALDEx2. The second assumption is that the amount of variation estimated by the statistical normalisation used by ALDEx2, the centred log-ratio normalisation, was lower than optimal. Nixon et al. [17] showed that the measured amount of variation was low for essentially all high-throughput sequencing datasets and that including some uncertainty was always optimal. As such, we included scale uncertainty of 0.5 standard deviations, which is the minimum recommended for this approach [21].

### 2.3 Vaginal metatranscriptomics captures the canonical community state types (CSTs)

Initial analysis of the London and Europe datasets recapitulated numerous prior studies on compositional differences between healthy and BV microbiomes. Hierarchical clustering of species-level metatranscriptome profiles exhibited clusters strongly reminiscent of the canonical CSTs first reported by Ravel and colleagues [11] (Figure 2A). We identified three *Lactobacillus*-dominant clusters (*L. crispatus*, *L. iners*, *L. jensenii* ), a cluster of anaerobic species at various relative abundances, and a final cluster characterised by a mix of *L. iners* and several BV-associated species.

**Fig 2.**
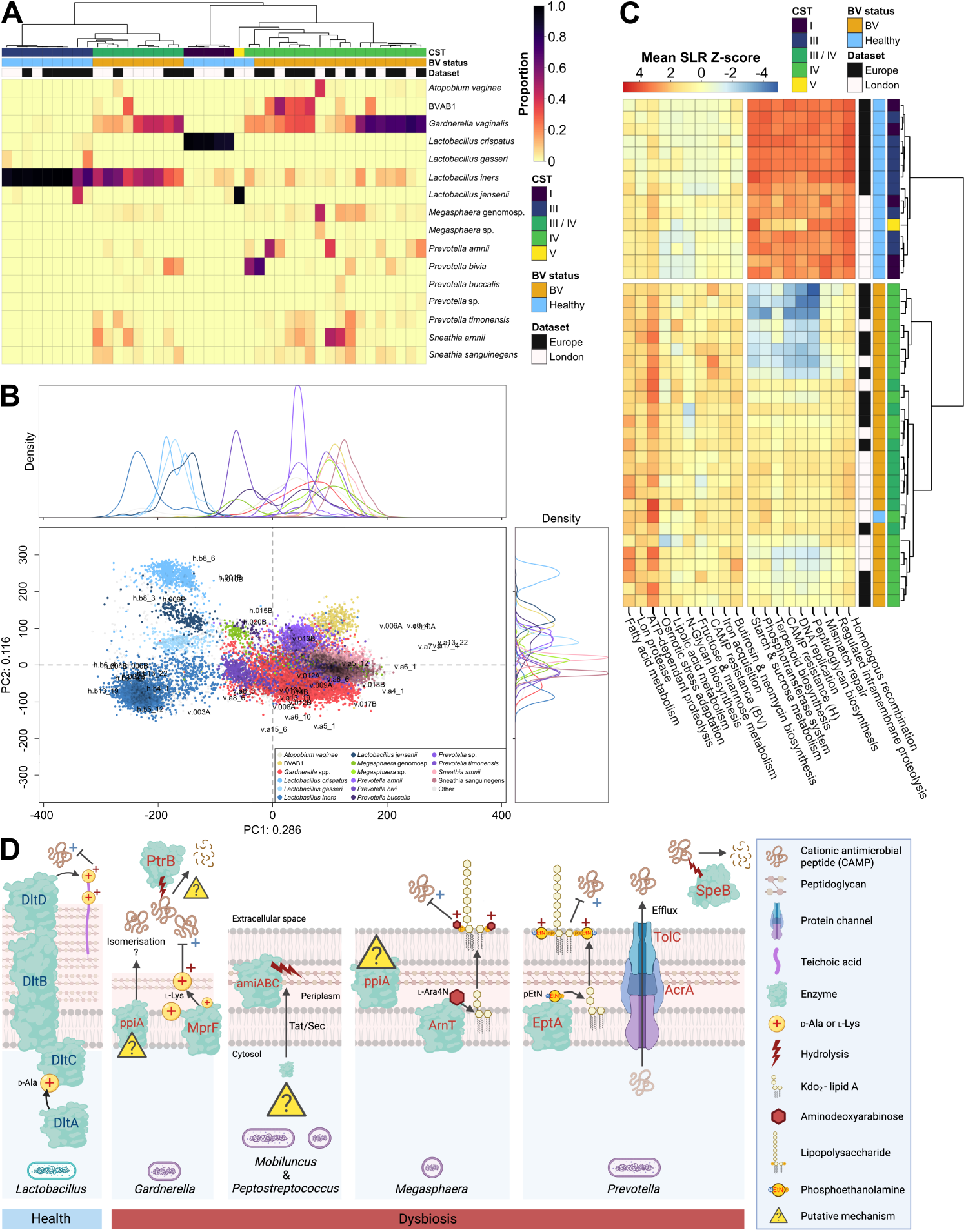
Health and BV-associated species of the vaginal microbiome employ differing strategies for host CAMP resistance: Vaginal microbiome composition (gene-level mapping, coloured by species) within London and Europe datasets was assessed by (**A**) hierarchical clustering of metatranscriptomes and (**B**) compositional principal component analysis (post-SRI); only species represented by ≥75 genes are shown. Marginal density plots show gene distribution by species across PC1 and PC2. (**C**) Gene assignments for each sample were grouped by KEGG orthology (KO) term regardless of species and median scaled log-ratio (SLR) values calculated by ALDEx2 were aggregated into a Z-score of the mean SLR value per KEGG pathway. Data for the top 10 differential pathways in both health and BV are plotted (all post-SRI absolute effect sizes and expected false discovery rates are >1 and <0.01, respectively). (**D**) Known and putative mechanisms of CAMP resistance within vaginal metatranscriptomes of the London/Europe datasets (created with BioRender).

Metatranscriptome reads that mapped to the gene and organism level of healthy and BV patients were readily observed to group by species on a compositional principal component biplot [30] (hereafter PCA plot). Health-associated *Lactobacillus* species were cleanly separated across the first principal component from the various anaerobic species associated with BV. Likewise, BV status was easily distinguished for all samples across PC1 (Figure 2B). We further observed that genes assigned to *Gardnerella vaginalis* and either phylogroup of *Megasphaera* did not form the discrete clusters observed in other species and were instead dispersed across PC1. While *Megasphaera* phylogroups have not been formally distinguished as individual taxa [31], the genus, *Gardnerella*, has recently been delineated into at least 13 genomic species [32], four of which have official standing in nomenclature (though they are is not presently included in the VIRGO database taxonomy).

### 2.4 Resistance mechanisms against host cationic antimicrobial peptides differ between healthy and BV metatranscriptomes

Differential abundance analysis between healthy and BV metatranscriptomes with ALDEx2 demonstrated significant differences in expression for a multitude of pathways, many of which reflect previously reported differences between the two conditions (Figure 2 & Supplementary Figure S2). For example, expression of genes involved in butanoate metabolism [33] and sialidase activity [34]– hallmarks of BV and some species of *Gardnerella*, respectively– were elevated among BV metatranscriptomes, while peptidoglycan biosynthesis genes were highly expressed among healthy samples dominated by Gram-positive lactobacilli.

One of the most striking newly observed differences between healthy vs. BV samples was observed in genes involved in resistance to host-encoded cationic antimicrobial peptides (CAMPs) (Figure 2C). Five KEGG orthology (KO) terms assigned to this pathway were defined as differential (four over-expressed in healthy samples, one over-expressed in BV samples) and absolute effect sizes for these KO terms were large (range 1.02 - 1.77). Differential CAMP resistance KO terms in healthy samples corresponded to all four genes of the DltABCD operon– a ubiquitous pathway among Gram-positive bacteria [35]– which enables d-alanylation of peptidoglycan-anchored teichoic acids, resulting in a reduction of the negative surface charge. These KO terms were initially annotated in KEGG as ‘*Staphylococcus aureus* infection’.

Among metatranscriptomes from the BV group, only a single KO term for CAMP resistance was defined as differentially abundant, corresponding to the periplasmic N-acetylmuramoyl-L-alanine amidases AmiABC, which are critical for cleaving peptidoglycan during cell division and separation–particularly under acidic conditions [36]. However, we were unable to find any reports corroborating a link between any of these amidases with CAMP resistance. Accordingly, we speculate that it may be a misassignment in KEGG which also reflects the higher proportion of Gram-negative species within BV microbiomes.

Given the importance of host-microbe interactions during microbial colonisation, we next investigated the expression of all genes present in the KEGG CAMP resistance pathway (map01503) among BV samples in the London and Europe datasets. A further eight genes in this pathway were also present in the London and Europe datasets (Figure 2D), with functions including those that decrease the net negative surface charge for repulsion of CAMPS, efflux of CAMPs, and direct CAMP hydrolysis; however, we were unable to find confirmatory studies linking two genes (PpiA/PtrB) to CAMP resistance. In multiple instances, a single genus was responsible for the majority of genes assigned to a given KO term.*Gardnerella* spp., for example, contributed 17 of the 34 genes with known taxonomy which were assigned to the phosphatidylglycerol lysyltransferase, MprF (KO14205; [37]), while 10 of 12 genes assigned to the lipid A ethanolaminephosphotransferase, EptA (K03760; [38]), originated from several species of *Prevotella*. In both cases, other species typically contributed only a single gene to both KO terms.

During our search for CAMP resistance genes, we also noted the presence of several genes assigned to the KO term, ‘African trypanosomiasis’ (K01354), which was unexpected considering the vaginal niche. This KO term represents the prolyl oligopeptidase, PtrB, which degrades peptides smaller than ∼30 amino acids via hydrolysis at the carboxyl side of basic amino acids [39]. A total of 58 genes in the London and Europe datasets were assigned to this KO term: 31 were of unknown taxonomy, 23 were assigned to *G. vaginalis*, 2 to *Bifidobacterium longum*, 1 to *Mobiluncus mulieris* and 1 to *Streptococcus pneumoniae*. Although we could not find literature supporting a role for PtrB in CAMP resistance, given the abundance of basic amino acids in CAMPs, we hypothesise that such oligopeptidase activity might represent an additional strategy used by BV-associated taxa for evading host immune responses.

The potential misassignment of multiple KO terms for CAMP resistance functions led us to screen the list of differential KOs for similarly suspicious functions, and confirm that KO terms with no assigned function were truly unknown. Two KO terms flagged due to an assignment of ‘Epithelial cell signaling in *Helicobacter pylori* infection’ corresponded to the acid-activated urea channel, UreI (K03191), and the putative U32-family collagenase, PrtC (K08303). Genes contributing to the latter function were abundant among several BV-associated taxa, including several species of *Prevotella* and *Sneathia*, *Atopobium vaginae*, and BVAB1. Both terms were over-represented among BV metatranscriptomes (Supplementary Figure S2) and may bear relevance to BV, given the involvement of UreI in pH resistance among clinically important human pathogens [40] and the identification of PrtC in several vaginal *Prevotella* and *Porphyromonas* and isolates capable of type I collagen degradation [41, 42]. Likewise three terms initially lacking functional information, were respectively found to correspond to the periplasmic iron-binding shuttle, FbpA (K02012), the non-heme ferritin iron storage protein, FtnA (K02217), and the acid-inducible, cytoplasmic membrane-bound iron transporter, EfeU (K07243). Iron acquisition is directly linked to colonisation and virulence in many important human pathogens [43]; therefore, the differences in iron requirements between BV-associated taxa and health-associated lactobacilli may present a therapeutic opportunity [44].

### 2.5 Subgroups of BV metatranscriptomes are mainly delineated by expression of genes involved in motility, chemotaxis, and co-factor biosynthesis

Hierarchical clustering of samples based on all differentially abundant pathways revealed a clear split among BV metatranscriptomes, suggesting the existence of at least two functional subgroups within the BV population (Supplementary Figure S2). A comparison of the two main BV subgroups at the functional level demonstrated that genes involved in motility (flagellar assembly), heme and cobalamin/vitamin B_12_ biosynthesis (porphyrin metabolism) and chemotaxis were the main drivers of this difference within the classical archetype of molecular BV (i.e. CST IV with/without BV symptoms; Figure 3, top). Based on the taxonomic composition of these samples, the presence or absence of BVAB1 (BV-associated bacterium-1; recently reclassified as “*Candidatus* Lachnocurva vaginae” [45]) appears to explain these differences almost entirely (Figure 3, bottom).

**Fig 3.**
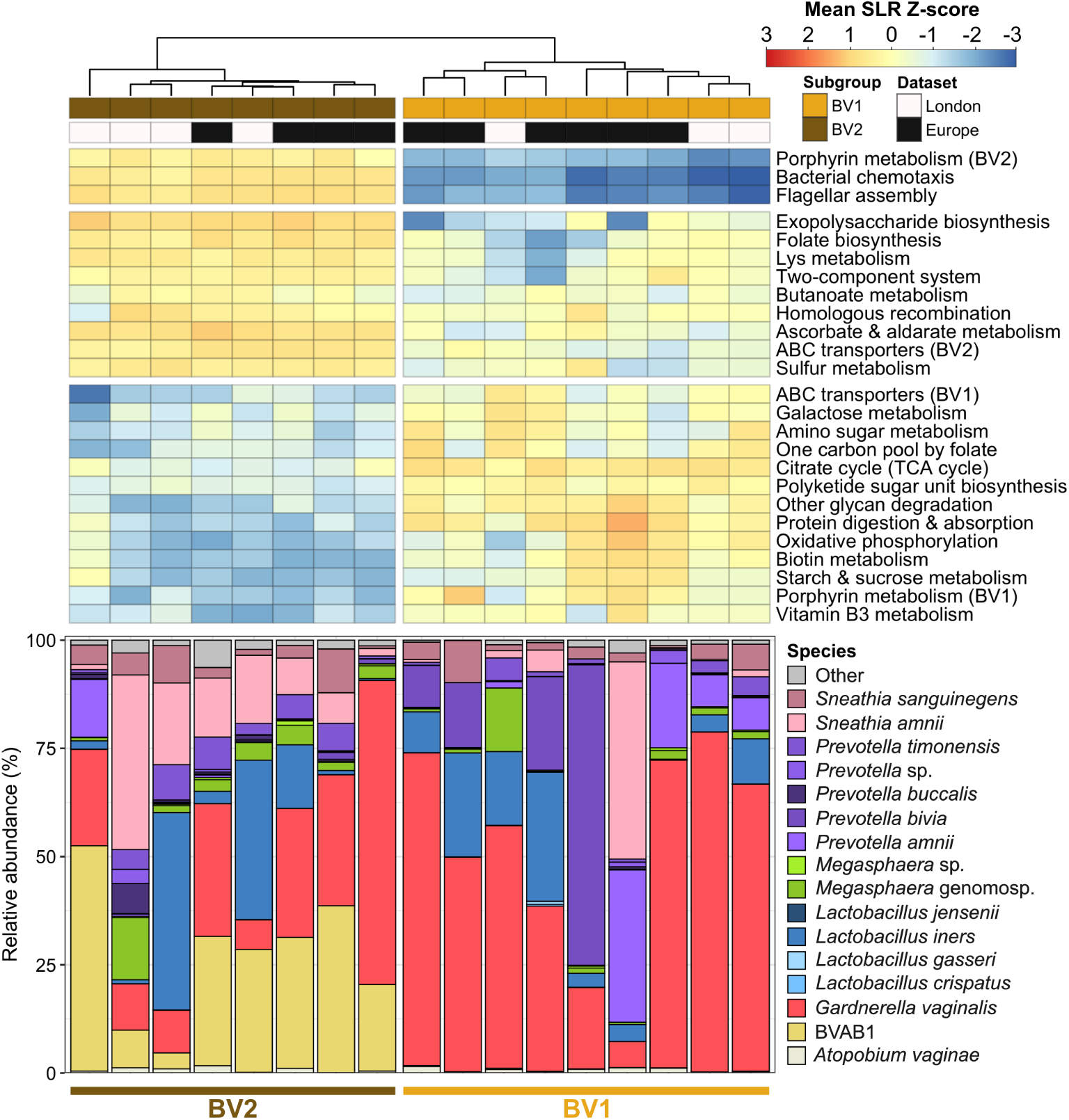
Differential expression of motility, chemotaxis and co-factor biosynthesis genes in distinct BV subgroups is largely due to the presence of BVAB1: The top 25 differentially abundant pathways among samples from the two main BV subgroups in London/Europe datasets are expressed as a Z-score calculated from the mean scaled log-ratio (SLR) values of all KO terms assigned to a given pathway (individual SLR values represent a median across 128 Monte-Carlo instances). All post-SRI absolute effect sizes and false discovery rates calculated by ALDEx2 were >1 and <0.01, respectively). Stacked bars below represent microbiome composition expressed as proportional abundance. Taxa represented by <75 genes are collapsed into ‘Other’.

When considering the taxonomy of reads associated with the KO terms for ‘bacterial chemotaxis’, reads originating from BVAB1 comprise the majority. The eight KO terms over-represented in BV2 in the London/Europe datasets corresponded to various genes within the Che signalling pathway [46], including the sensor histidine kinase (CheA), signal transducers (CheW/V), regulatory methyltransferases/demethylases (CheR/B), and methyl-accepting chemoreceptors (mcp). The KO term for the Che response regulator (CheY; activates the flagellar motor switch upon phosphorylation by CheA) was detected at higher proportions in BV2 samples than in BV1 samples, but this difference was marginally below our effect size threshold of significance (effect = -0.897).

Likewise, reads assigned to the KO terms for ‘flagellar assembly’ mostly originated from BVAB1, as well as *Lactobacillus iners*, *Mobiluncus curtisii* and *Mobiluncus mulieris*. A variety of flagellar components were represented by these KO terms [47], including the major flagellar filament subunit (fliC), rotary protein (motA), hook proteins (flgE/K/L and fliD/K), motor switch proteins (fliM/G), regulatory factors (flgM and flaG), type III protein export proteins (flhA and fliH/I), and C- and MS-ring proteins (fliF/N).

The presence of a KO term corresponding to ‘exopolysaccharide biosynthesis’ among the differential pathways was particularly notable, given its function as poly-*β*- 1,6-acetyl-d-glucosamine synthase (PgaC/IcaA; K11936 [48]). This polysaccharide is a key component of biofilm extracellular polymeric substance produced by both Gram-negative and Gram-positive species, and has been implicated in initial cell-surface attachment and later lateral agglomeration during biofilm development and expansion [49]. Biofilm involvement is a prominent feature of BV [50] and vaginal epithelial cells with adherent biofilms (clue cells) are a diagnostic feature during microscopic analysis of Gram-stained, wet-mount vaginal smears [51]. Once more, reads belonging to this KO term were strongly associated with BVAB1 (and to a far lesser extent, *Prevotella amnii* ).

Reads corresponding to the KO terms for ‘porphyrin metabolism’ showed no association with BVAB1. Of the three KO terms over-represented in the BV2 subgroup, almost all reads assigned to the cobaltochelatase, CobN, originated from *Prevotella timonensis*, while the majority of reads assigned to the cobalamin adenosyltransferase, PduO, and the porphobillinogen synthase, HemB, were of unknown taxonomy. The KO term elevated in BV1 metatranscriptomes corresponds to uroporphyrinogen III synthase (hemD), and reads assigned to this term were largely contributed by *Prevotella bivia* and *Prevotella amnii*. HemB/D are involved in both heme biosynthesis via *δ*-aminolevulinate and *de novo* production of the biologically-active (co-factor) form of vitamin B_12_, adenosylcobalamin. PduO plays a role in the cobalamin salvage pathway, converting exogenous cobamide taken up via the BtuBCDF transport system into adenosylcobamide, while CobN oversees the incorporation of cobalt into the tetrapyrrole ring of hydrogenobyrinic acid a,c diamide as part of the *de novo* biosynthesis of adenosylcobalamin [52].

The existence of distinct subgroups of BV– particularly those delineated by expression of genes involved in motility and biofilm formation– led us to speculate that symptomatic vs. asymptomatic BV may be linked to one subgroup or the other. Unfortunately, clinical metadata for the London/Europe datasets was limited to Nugent scores and vaginal pH, which precluded this analysis. However, extensive self-reported data on BV symptoms and vaginal pH was collected for the Virginia dataset as part of the MOMS-PI study on pre-term birth [7]. Therefore, we sought to replicate all of the above observations in the Virginia dataset and determine if there was any association of BV symptoms with any identified subgroups.

### 2.6 Pregnancy-related compositional changes within vaginal metatranscriptomes and lack of functional correlation with pre-term birth

Prior to validating any previous observations, we investigated if pregnancy introduced major taxonomic shifts which could complicate comparisons between London/Europe (non-pregnant) and Virginia (pregnant) datasets. We determined the CSTs present within the Virginia dataset and proceeded to define ‘molecular’ states of health and BV based on hierarchical clustering of metatranscriptomes at the functional (KO) level (Supplementary Figure S3). Virginia metatranscriptomes separated into two clear groups– the first, characterised by higher vaginal pH, a preponderance of CST IV samples and the absence of CST I samples, was taken to represent molecular BV. The other largely contained samples belonging to CSTs I, III, and V and was defined as molecular health. Notably, there appeared to be no pattern linking CST or molecular heath/BV with self-reported abnormal vaginal discharge, odor, or irritation.

After processing the three datasets together, we compared the post-SRI SLR values returned by ALDEx2 for all features from the species represented by at least 75 genes, stratified by dataset and BV status (Figure 4A). *Lactobacillus* depletion and anaerobic overgrowth in BV metatranscriptomes of all datasets was evident, and the effect of recent antimicrobial therapy for symptomatic BV within the Europe dataset was clear, manifesting as lower abundances of non-*L. iners* lactobacilli and higher abundances of BV-associated taxa even among healthy samples. However the most striking differences between pregnancy states lay in the abundances of *Sneathia amnii*, *Sneathia sanguinegens*, and *Gardnerella vaginalis* in BV samples. Abundances of the two *Sneathia* species were lower in the Virginia dataset than in London or Europe– particularly *S. sanguinegens*– with a larger proportion of samples with extremely low abundances of reads corresponding to these taxa. Concomitantly, *Gardnerella vaginalis* abundances and to a lesser degree, those of *Atopobium vaginae*, were higher within the Virginia dataset than either non-pregnant dataset.

**Fig 4.**
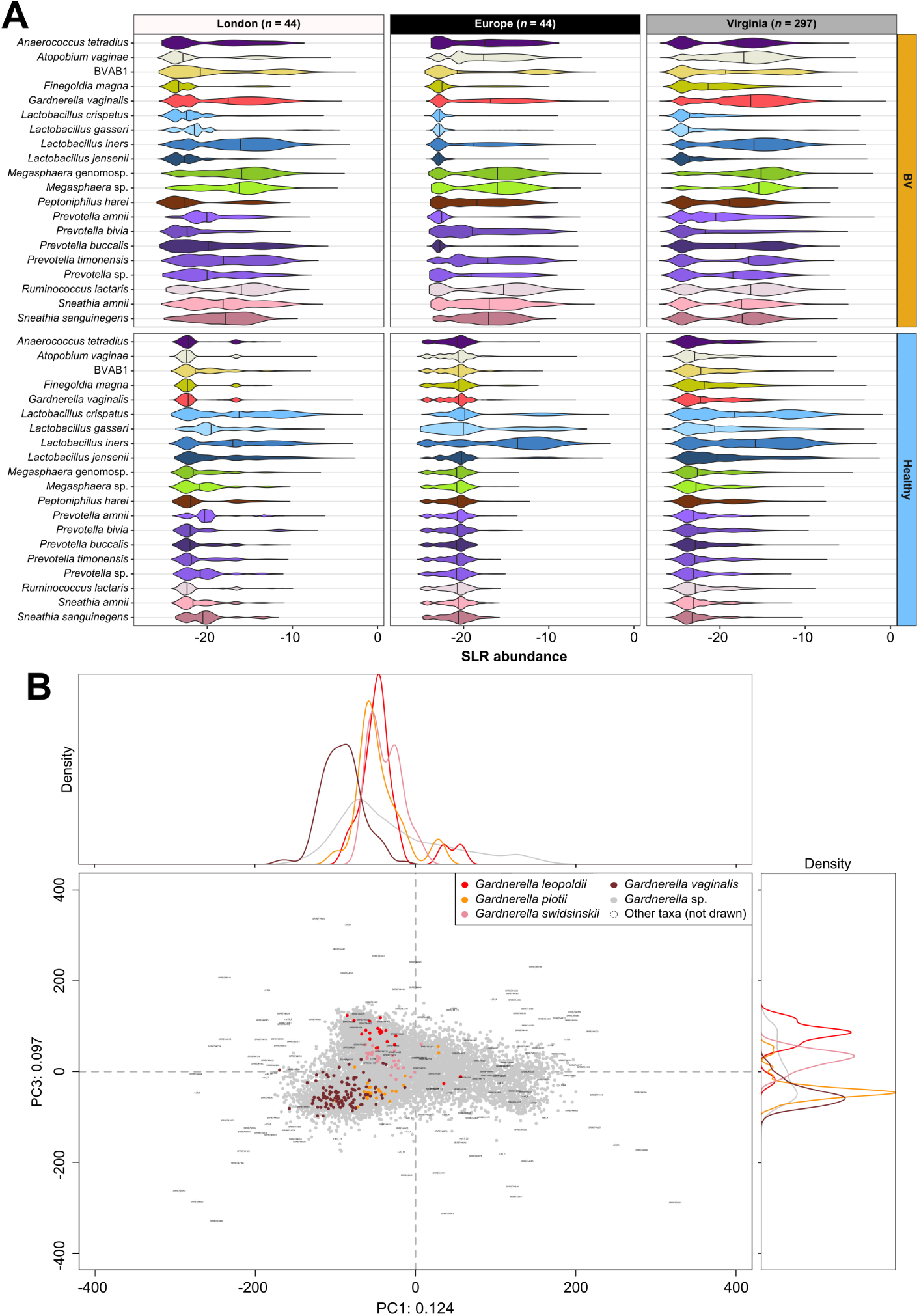
Subtle changes in vaginal microbiome composition and delineation of *Gardnerella* species within the Virginia validation dataset: Vaginal metatranscriptomes from all datasets in this study underwent filtering and scaled log-ratio (SLR) transformation with ALDEx2. (**A**) SLR values for all genes grouped by species were visualised as violin plots, stratified by dataset and BV status. Vertical lines on violins indicate taxa medians. (**B**) Genes unique to each of the four *Gardnerella* species with standing in nomenclature were identified and highlighted on a PCA plot of the same SLR-transformed data used in (A). Genes from all other species are not visible, but were included in the ordination. Density plots show gene distribution across PCs.

### 2.7 Discrete clusters of reads from named *Gardnerella* species are detectable within metatranscriptomes of all datasets

The prevalence of *Gardnerella vaginalis* in the Virginia dataset prompted us to ask whether the four named *Gardnerella* species with current standing in nomenclature could be identified and delineated within the broadly distributed group of reads assigned to the genus, *Gardnerella*, in the VIRGO database. Using publicly available, complete genome sequences (see methods), we identified genes unique to *G. leopoldii*, *G. piotii*, *G. swidsinskii* and *G. vaginalis* with Paneroo [53] and aligned sequences in the combined London/Europe/Virginia dataset which were labelled as *Gardnerella vaginalis* to these genes. Re-labelling the gene taxonomy according to these BLAST hits (conservatively filtered by E-value and alignment coverage), we then visualised reads assigned to the genus, *Gardnerella*, within the ordination of all features in the combined dataset (Figure 4B).

Separation of named *Gardnerella* species was seen across all features assigned to the genus, particularly those belonging to *G. vaginalis* and *G. leopoldii*. Similar separation at the species level was also seen when considering the London/Europe and Virginia datasets, separately (Supplementary Figure S4). Curiously, we observed an absence of genes unique to the named species for samples with positive PC1 scores, despite the discrete clustering of several hundred *Gardnerella* features. Such clusters may represent the multiple, unnamed genome species of *Gardnerella*, for which there are a paucity of representative complete genome sequences.

### 2.8 Key findings on from discovery datasets can be replicated in the validation dataset

Having shown that the vaginal metatranscriptome datasets were comparable despite differing pregnancy status, we asked whether the discoveries made in the London/Europe datasets could be replicated in the Virginia dataset.

Using the previously defined groups of molecular health and BV for SLR-transformation and data asymmetry correction, we visualised functional metatranscriptomes via PCA (Supplementary Figure S5). Following scale simulation with the same parameters as before, typical housekeeping KO terms corresponding to ribosomal translation machinery or aminoacyl-tRNA biosynthesis were again centred around the location of no difference. Metatranscriptomes clearly separated across PC1 based on molecular health vs. molecular BV status; however, we also observed an evident split within the molecular health group, driven by dominance of either *L. iners* (health subgroup 1) or *L. crispatus* (health subgroup 2). For consistency with earlier analyses, subgroups of molecular health were treated as a single group for comparison with molecular BV.

Formal differential abundance analysis mirrored the results obtained from the London/Europe datasets. CAMP resistance was once again among the most differential pathways, with the DltABCD operon highly expressed among healthy metatranscriptomes (Figure5A). All CAMP resistance KOs summarised in Figure 2D were also present in the Virginia dataset, and the same species-specific associations were observed (e.g. MprF with *Gardnerella vaginalis*). Several additional KOs belonging to the KEGG ‘CAMP resistance’ pathway were also detected, including several two-component regulatory systems (PhoPQ, CpxAR, BasRS), although these were represented by *<*100 reads each (likely a result of the greater sequencing depth in this dataset).

Almost all KO terms over-represented among BV metatranscriptomes in the discovery dataset were also differentially expressed in the validation dataset. KOs corresponding to butanoate metabolism, the U32-family putative type I collagenase (PrtC), the iron transport and storage proteins (EfeU, FbpA, FtnA), but not the acid-activated urea channel (UreI) exhibited significantly higher expression among BV metatranscriptomes. Unlike the London/Europe dataset, the single CAMP resistance KO term over-represented among BV metatranscriptomes corresponded to the phosphatidylglycerol lysyltransferase, MprF (absolute effect size = -1.23), rather than the N-acetylmuramoyl-L-alanine amidases, AmiABC (Supplementary Figure S6). Assessment of the signature functions defining health and BV groups overlapped well with differential abundance analyses and discriminated these groups with high accuracy (AUC = 0.997, Supplementary Figure S7A-C):CAMP resistance was a key functional signature of healthy metatranscriptomes, whereas butanoate metabolism and TCA cycle genes were among the characteristic features of BV.

As in the discovery dataset, the initial clustering of Virginia metatranscriptomes at the functional level revealed multiple sub-populations of BV metatranscriptomes. Focusing on the two largest subgroups (BV1, *n* = 57; BV2, *n* = 56), we tested whether the same differences between BV subgroups identified in the London/Europe dataset could be observed among Virginia BV samples. Differences in expression of genes involved in flagellar assembly, chemotaxis, and exopolysaccharide biosynthesis were again the among the largest drivers of subgroup separation (Figure 5B), with flagellar assembly identified as the sole contributor to the functional signature of one subgroup (AUC = 0.994, Supplementary Figure S7D-F). All KO terms from flagellar assembly and chemotaxis pathways identified as differential within the discovery dataset were again differentially expressed between BV subgroups in the validation dataset. Additionally, several KO terms from these pathways not identified as differential previously were among the differentially expressed KOs in the validation dataset, including the flagellar basal body rod protein, FlgD (K2389), the sigma factor regulating transcription of flagellar genes, FliA (K02405), the chemotaxis response regulator that interacts with the flagellar motor switch, CheY (K03413), and the heme-based aerotactic oxygen sensor, HemA/T (K06595). Exopolysaccharide biosynthesis was represented by a singular KO term (K11936) corresponding to the same poly-*β*-1,6-acetyl-d-glucosamine synthase identified in the discovery dataset. The only major difference between BV subgroup analyses in the discovery and validation datasets concerned porphyrin metabolism. Only a single KO term from this pathway was differentially expressed between subgroups: the porphobillinogen synthase, HemB.

**Fig 5.**
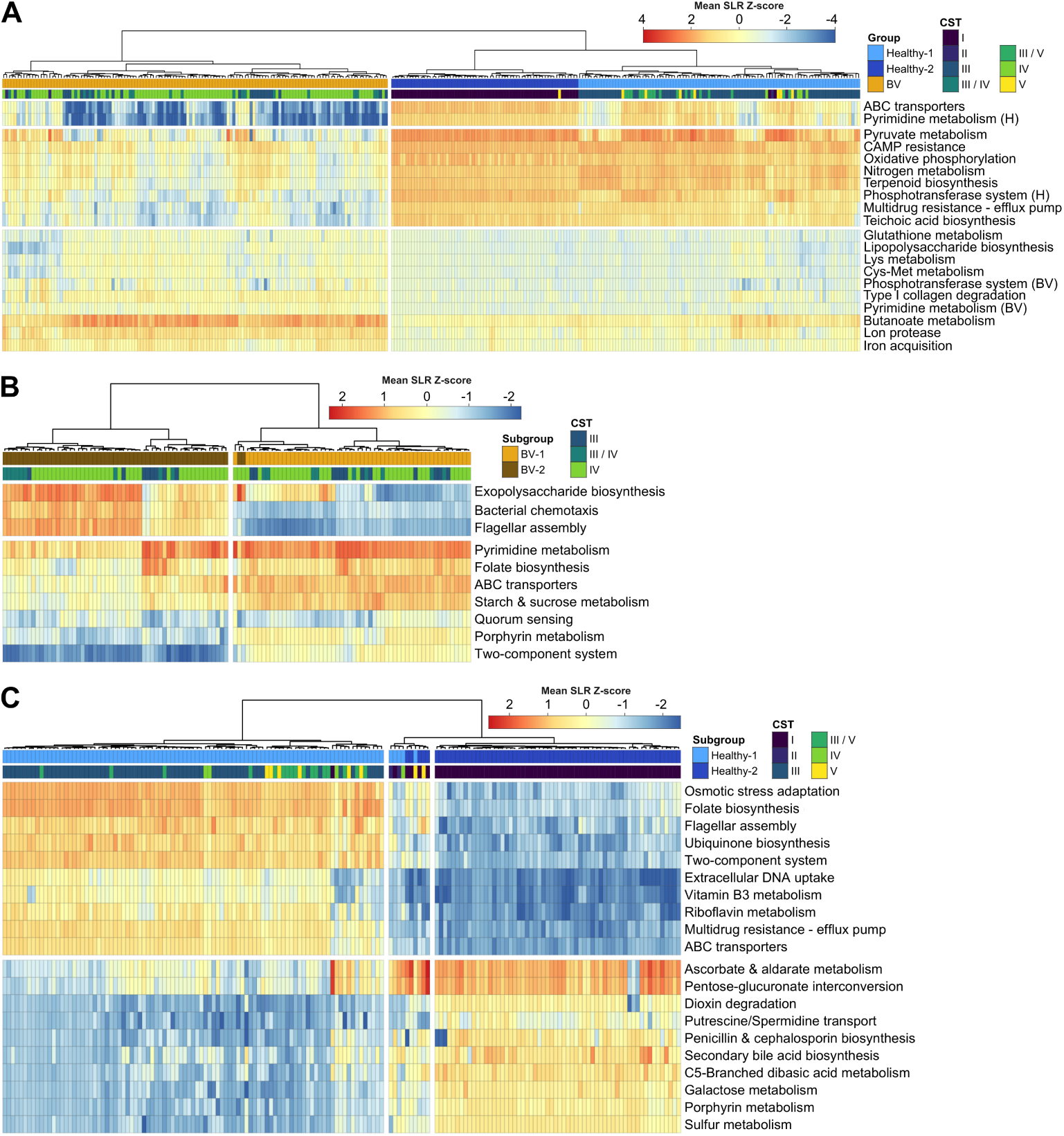
Prior observations on CAMP resistance and BV subgroups can be replicated in a large-scale validation dataset: Filtered metatranscriptome feature tables underwent scaled log-ratio (SLR) transformation with ALDEx2. Gene assignments for each sample were grouped by KEGG orthology (KO) term regardless of species and median SLR values from ALDEx2 were aggregated into a Z-score of the mean SLR value per KEGG pathway. These data were plotted for three comparisons of Virginia dataset samples: (**A**) molecular health vs. molecular BV; (**B**) molecular BV subgroups; (**C**) molecular health subgroups. For (**A**) and (**C**), the top 10 most differential pathways for health and BV are shown. All absolute effect sizes and expected false-discovery rates are ≥1, <1%, respectively.

All previous species-specific associations were recapitulated among BV sub-groups: BVAB1/*L. iners*/*Mobiluncus* spp. with flagellar assembly and chemotaxis, and BVAB1/*P.amnii* with exopolysaccharide biosynthesis. In fact, the presence of BVAB1 in BV1 but not BV2 samples again appeared to be responsible for a large degree of the separation of these subgroups (Supplementary Figure S8). We did however note that the expression of flagellar genes between BV1 and BV2 groups appeared to be strain-specific, such that the difference in the total number of reads which were assigned to this pathway and originated from *L. iners* spanned three orders of magnitude (1,211,315 reads vs. 2,344 reads). Critically, we were able to replicate all of the above findings when genes were aggregated by function using the eggNOG classification system (Supplementary Figure S9). EggNOG terms corresponding to flagellar components, chemotactic signalling machinery and exopolysaccharide biosynthesis/biofilm formation were among the most differential functions when comparing the same BV subgroups.

Finally, the size of the validation cohort allowed us to investigate the clear separation among metatranscriptomes within the molecular health group which appeared to be due to dominance by either *L. crispatus* or *L. iners*. Differentially expressed functions generally reflected metabolic differences between these two taxa (Figure 5C & Supplementary Figure S10). Of interest, expression levels of genes involved in flagellar assembly among *L. iners*-dominant samples were significantly higher, while those encoding a putative polyamine transport system specific for putrescine and spermidine were among the most differentially expressed transcripts for *L. crispatus*-dominant samples. Previous work has demonstrated that vaginal isolates of *L. crispatus* are capable of producing or degrading biogenic amines, including putrescine, depending on the strain [54]. Moreover, this transport system was also detected in a recent RNA-seq study of the vaginal environment [20]. Equally of note was a single KO term assigned to ‘dioxin degradation’ (K01821; PraC/XylH); of all reads in the validation dataset assigned to this KO term, 99.9% originated from lactobacilli (74.6% from *L. crispatus*). That dioxins are classed as persistent chemical pollutants generated as industrial by-products makes it highly unlikely that they are truly metabolised within the vaginal environment [55]. We note that this KO term corresponds to 2-hydroxymuconate tautomerase activity and that 2-hydroxymuconate and catechol are the products of enzymatic reactions involving enzymes of the EC class 3.7.1 (hydrolases acting on C-C bonds in ketonic substances). Furthermore, 2-hydroxymuconate can be metabolised to pyruvate via *γ*-oxalocrotonate and 4-hydroxy-2-oxopentanoate. Though entirely speculative, this could theoretically represent a mechanism by which vaginal estrogen may be used as an energy source by *L. crispatus*.

## 3 Discussion

Overall, our meta-analysis of three independent vaginal metatranscriptome datasets identified differing mechanisms by which species associated with health and BV evade local host immune responses and persist within the vaginal niche. We also observed distinct functional subgroups within the BV population of all datasets, largely driven by the presence of BVAB1 and differing in expression of pathways potentially of relevance to BV pathogenesis, including motility and biofilm formation. Critically, these findings were robust to the different methodological approaches employed by the original study authors, including the large discrepancy between study population demographics, as well as different functional classification systems employed in our analyses. Our findings, together with other known host-microbe interactions in the vaginal environment, are summarised in Figure 6.

**Fig 6.**
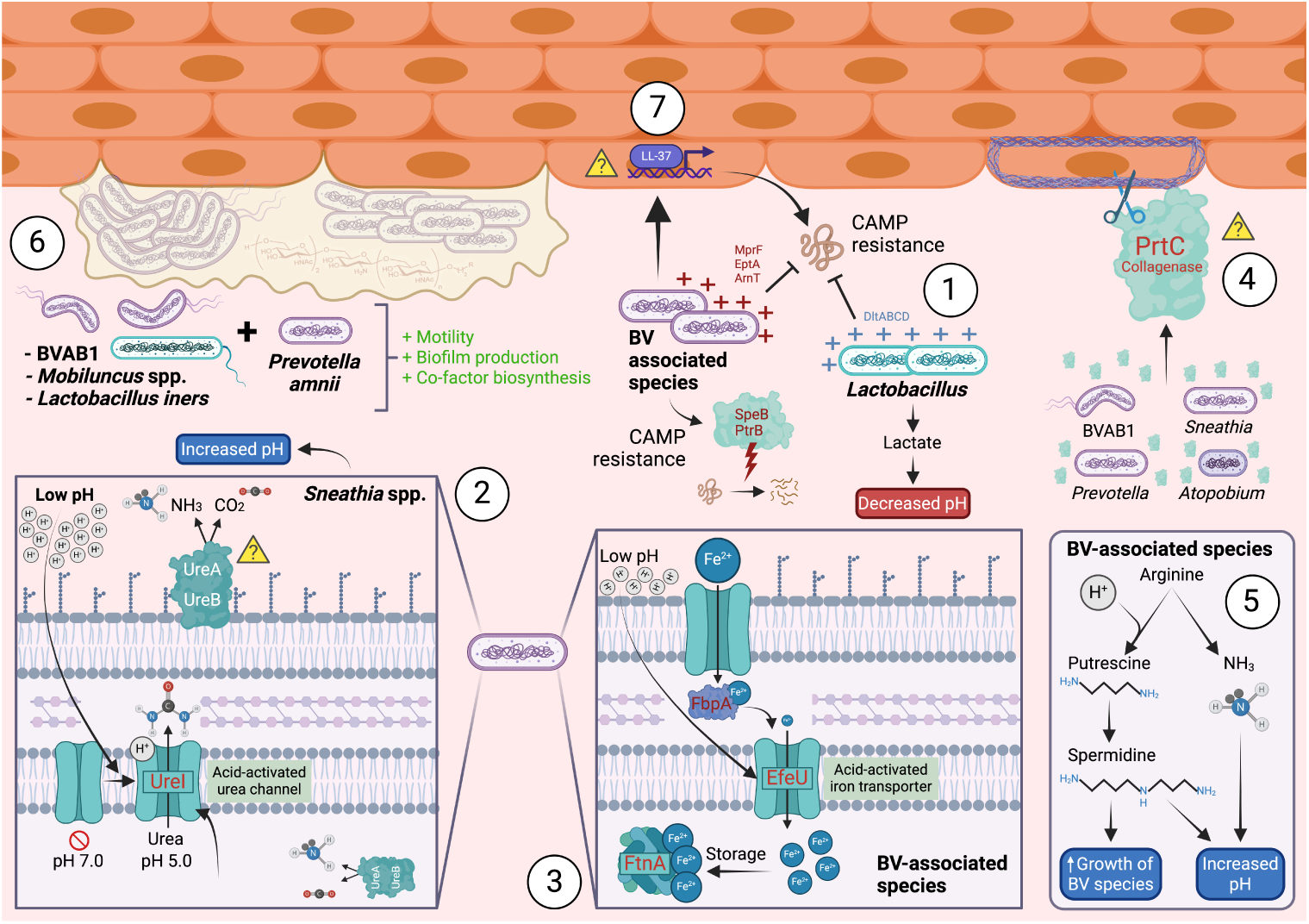
Summary of microbial persistence mechanisms and host-microbe interactions in the vaginal environment: All functions assigned according to the Kyoto Encyclopedia of Genes and Genomes. Yellow triangles indicate putative mechanisms. **1.** Resistance to host cationic antimicrobial peptides (CAMPs; see Figure 2D) involves reducing net negative surface charge or direct CAMP degradation. **2.** Low vaginal pH triggers the opening of the inner membrane acid-activated urea channel, UreI (K03191). Cytoplasmic and/or surface-adsorbed UreA/B urease may then serve to raise local pH. **3.** Low vaginal pH also triggers opening of the acid-activated periplasmic iron channel, EfeU, permitting transport of iron across the periplasm by FbpA (K02012) and its intracellular accumulation by the non-heme ferritin, FtnA (K02217). **4.** Production of a U32-family peptidase orthologous to PrtC (K08303) found in vaginal *Porphyromonas* isolates may degrade collagen in the extracellular matrix of the vaginal mucosa. **5.** Production of the biogenic amines, putrescine and spermidine, from arginine. These polyamines, along with the liberation of ammonia during their synthesis, raise vaginal pH and can promote the growth of BV-associated taxa. **6.** Subgroups of BV are delineated by biofilm formation, motility and co-factor biosynthesis; BVAB1 and *Prevotella amnii* synthesise poly-*β*-1,6-acetyl-d-glucosamine via PgaC (K11936). BVAB1, *Mobiluncus* spp., and *L. iners* express various flagellar and chemotaxis genes as part of the ‘motile’ BV genotype. **7.** Induction of LL-37 production and secretion by *Gardnerella* spp., as occurs at the urothelium during urinary tract infections. Similar mechanisms may be employed by *Gardnerella* spp. in the vagina to out-compete lactobacilli or other taxa. Created with BioRender.

The adoption of the principles of SRI into the Bayesian models constructed by ALDEx2 was critical for analysis of these vaginal metatranscriptome datasets. The original ALDEx2 algorithm acknowledged that sequencing datasets represent a single point estimate of the sampled environment which does not reflect the true community composition. ALDEx2 generates a posterior distribution of the observed data by repeated sampling from a multinomial Dirichlet distribution [23]. This posterior distribution undergoes a centred log-ratio normalisation using the geometric mean (*G_n_*) of all features in a given sample. Recently, Nixon *et al.* [17] demonstrated that in doing so, ALDEx2 makes an inherent assumption about the scale of the community; that the inverse of the geometric mean of the observed counts is exactly equal to the scale. However, the estimate of the geometric mean calculated by ALDEx2 is, by definition, a precise but inaccurate measure of scale since all information about community scale is lost during the sequencing process. Thus, even small errors in the scale assumption can have a drastic effect on estimates of the true log fold-change of a given feature between conditions. For this reason, these authors recommended that models should account for errors in the scale assumption during normalisation to prevent converging on a precise but inaccurate estimate of scale and hence differential abundance.

We identified several genes involved in CAMP resistance among the most differentially expressed pathways between healthy and BV metatranscriptomes. Crucially, this finding was robust to replication across three independent datasets which were distinct in terms of geography,sampling methodology, pregnancy status and ethnicity. Given the well-reported effect of ethnicity on vaginal microbiome composition [11, 56], the replication of almost all findings across exploratory and validation datasets is remarkable and underscores the value of correct data normalisation. In addition to uncovering several mechanisms by which health– and BV-associated species persist within the vaginal niche, this finding also sheds light on important co-evolutionary processes within the vaginal microbiome. Kiattiburut *et al.* [57] demonstrated how BV-associated species implicated in urogenital infections (including *G. vaginalis*; W. Kiattiburut & J. Burton, personal communication) are capable of inducing the production and secretion of host CAMPs such as LL-37 by urothelial cells. As lactobacilli do not evoke the same response [57], it is feasible that this represents a unique strategy employed by *Gardnerella* to redirect host LL-37 towards lactobacilli. That both taxa– along with a multitude of other vaginal microbiome constituents– express CAMP resistance genes points to ongoing co-evolution within these species and also explains earlier observations regarding the general resistance of vaginal lactobacilli to CAMPs [58]. Given the importance of *Gardnerella* species in the context of BV, targeting l-lysylation of membrane phospholipids by MprF and increasing susceptibility to CAMP-mediated lysis may prove to be of benefit when considering novel therapeutics for BV.

Many surveys of the vaginal microbiome employing amplicon sequencing have attempted to define subtypes of BV-associated CST IV, largely based around what was at the time thought to be subgroups/clades of *G. vaginalis* and other BV taxa [59, 60]. Our findings regarding functional subgroups of metatranscriptomes suggest that taxonomic classification using canonical CSTs is largely redundant– especially amongst BV populations. While BVAB1 abundance appeared to differ between BV subgroups, the abundance of other species contributing to differences in expression of flagellar (*L. iners*) or exopolysaccharide biosynthesis (*P. amnii* ) genes did not. For these species, differences between subgroups were likely due to strain-specific differences in gene content, as was the case in older culture-based studies reporting the isolation of flagellated and non-flagellated lactobacilli from vaginal swabs [61]. Given the importance of polymicrobial biofilms in BV pathogenesis (and more generally in resistance to antimicrobial agents), the existence of a motile, biofilm-forming BV sub-type may be of importance when considering the triggers for the onset of active BV symptoms and when investigating causes of treatment failure.

Differences in the expression of genes involved in iron uptake and storage is not particularly surprising, given that lactobacilli are not dependent on iron for growth.

Although sensitivity to iron levels may depend on nucleotide availability in some lactobacilli [62], most others are unaffected and instead rely on manganese and cobalt as co-factors in their essential redox enzymes [44]. It is noteworthy then that some BV subgroups express enzymes involved in cobalamin biosynthesis which, by definition, requires uptake of cobamide or other forms of cobalt. This may represent a strategy employed by BV-associated species to out-compete lactobacilli through the sequestration of a valuable resource. Likewise, expression of pH-dependent urea channels at the higher levels noted among BV metatranscriptomes compared to their healthy counterparts could reflect attempts by BV-associated taxa to counteract the low vaginal pH. Although detected at low levels, both urease subunits were also detected in our meta-analysis; this enzyme is central to similar pH-raising mechanisms that underpin virulence in pathogens such as *Helicobacter pylori* [40].

Finally, the identification of a collagenase present in several BV-associated species raises questions about the role played by collagen degradation BV pathogenesis, particularly regarding the cause vs. effect of dysbiosis. Several reviews [63, 64] point to the production of “collagenases and fibrinolysins” by various species of *Sneathia* and *Prevotella*, citing the work of Al-Mushrif *et al.*; however this work investigated the *in vitro* effects of organic acids on monocytes and only stated the fact that several pathogenic *Prevotella* produce these enzymes [65]. In the chain of papers cited, Hofstad ultimately demonstrated in 1984 the collagen- and fibrin-degrading activities of various *Prevotella* species (then classified as *Bacteroides*), though vaginal *Prevotella* were not studied [66]. More recently, preliminary work by Lithgow and colleagues confirmed that vaginal isolates of *Porphyromonas uenonis*, *Porphyromonas asaccharolytica Prevotella bivia* and *Sneathia amnii* all possess secretory collagenase activity [42]. While it remains to be experimentally proven how these enzymes access collagen within the extracellular matrix of the vaginal mucosa, damage to the epithelial barrier could favour bacterial attachment and influence biofilm formation, as is the case in the respiratory tract [67, 68]. Collagenase expression may also contribute to the production of the detached ‘clue cells’ [51] observed during clinical diagnosis, as these biofilm-covered epithelial cells could be released upon degradation of the extracellular matrix.

## 4 Conclusion

Overall, we have demonstrated the advantages of incorporating tools such as SRI into ‘omics analyses, showing how it can be used to identify differentially expressed functions which are robust to replication even across independent and highly divergent datasets. Our application of these tools to vaginal metatranscriptomic datasets opens several avenues of investigation in relation to potential therapeutic targets for BV which require further investigation to confirm their involvement in pathogenesis. The inclusion of SRI into the ALDEx2 platform should be of general use when examining multiple other metatranscriptomes and other problematic high throughput sequencing datasets.

## 5 Methods

### 5.1 Meta-analysis datasets

Three previously published vaginal metatranscriptome datasets were included in our meta-analysis: Macklaim *et al.* (London [24]), Deng *et al.* (European [25]), and Fettweiss *et al.* (Virginia [7]) (see ”Data availability and code” section for access information).

The London dataset was produced from vaginal swabs of the posterior fornix and mid-vaginal wall, collected from 22 women in 2013. Total RNA was extracted from swab fluid using a standard workflow [24] and RNA libraries were sequenced on an Illumina HiSeq 2000 instrument.

The European dataset was produced from vaginal lavage samples (2 mL saline) collected from 22 women between April 2014 and September 2015. Total RNA was extracted from 1 mL of vaginal fluid suspension using a Mo Bio PowerMicrobiome RNA Isolation kit and rRNA was depleted using a Ribo-Zero Gold rRNA Removal kit [25]. Paired-end RNA libraries were constructed with an Illumina ScriptSeq kit (2 x 110 bp reads) and sequenced on a HiSeq 2500 instrument.

The Virginia dataset was produced from vaginal swabs of the mid-vaginal wall, collected from pregnant women as part of the MOMS-PI study. Total RNA was extracted using a Mo Bio PowerMicrobiome RNA Isolation kit and rRNA was depleted using an Epicentre/ Illumina Ribo-Zero Magnetic Epidemiology Kit [7]. Paired-end RNA libraries were constructed with a KAPA Biosystems KAPA RNA HyperPrep Kit (2 x 150 bp reads) and sequenced on an Illumina HiSeq4000 instrument.

### 5.2 Re-processing of raw data

All FASTQ files were re-processed in the same manner to minimise differences across datasets due to bioinformatic pipelines. Raw reads were trimmed for quality (minimum Q20 across a 4bp sliding window) and length (cropped to 75 bp) using Trimmomatic (v0.39). Trimmed reads were then sequentially mapped to two different human genome assemblies (GRCh38 and T2T-CHM13) and all mapped reads were discarded. The remaining non-human reads were then mapped to the SILVA rRNA database (release 138) and all putative rRNA reads were discarded. All mapping steps were performed using Bowtie2 (v2.5.1) with the high accuracy option. Finally, non-human, rRNA-filtered reads were mapped to the VIRGO database–a non-redundant catalogue of genes from the human vagina [19] for taxonomic and functional assignment, and feature tables for each dataset were constructed separately, using scripts provided by the maintainers of VIRGO. Data were generated for gene-level, eggNOG [69] and KEGG [70] level analyses.

### 5.3 Filtering and batch-correction of metatranscriptomes

Feature tables from VIRGO were merged together in R and filtered with the CoDaSeq package to retain only the genes present in ≥30 % of samples, with a relative abundance of ≥0.005 % in at least one sample. The London and European datasets then underwent batch correction using the ComBat seq function from the sva package [28], prior to all downstream analysis in R. Batch correction was not performed for the Virginia dataset owing to its use as an independent confirmatory dataset. Analyses comparing metatranscriptomes by pregnancy status were performed on a merged feature of the London, Europe and Virginia datasets following filtering and batch-correction as above.

### 5.4 Species-level summaries of vaginal metatranscriptomes

Vaginal microbiome composition at the species level was summarised in terms of relative abundance (calculated from filtered, batch-corrected feature tables) using the pheatmap R package.

For visualisation of community composition via principal component analysis (PCA), batch-corrected feature tables underwent scaled log-ratio (SLR) transformation using the updated ALDEx2 package in R, prior to PCA with the prcomp function. The resultant compositional biplots were visualised using the CoDaSeq R package (https://github.com/ggloor/CoDaSeq). Gene functions were assigned using KEGG orthology (KO) numbers, eggNOG group membership and EC classification, as provided by VIRGO [19].

To differentiate the named species of the genus, *Gardnerella*, we obtained genomes of the species type strain, plus an additional (complete) genome, from NCBI for each of the following species (strains indicated in parentheses): *G. leopoldii* (UGent06.41; UMB6774), *G. piotii* (UGent18.01; JNFY15), *G. swidsinskii* (GS9838-1; JNFY3), and *G. vaginalis* (ATCC14018; NR001). Genomes were annotated for each species separately using prokka (v 1.13) [71], running with the ‘–proteins’ flag (using a .gff file for each species, pre-annotated by the NCBI PGAP tool). Output .gff files were used as input for pangenome analysis with panaroo (v1.3.3) [53] and the resulting presence-absence matrix was used to identify genes unique to each named *Gardnerella* species. Sequences corresponding to these ‘species marker’ genes were extracted from the VIRGO catalogue and aligned to a custom BLAST database consisting of VIRGO genes present in our merged dataset. BLAST results were filtered to retain only hits with ≥98 % sequence identity and the corresponding VIRGO genes were used to estimate the location of the four named *Gardnerella* species on PCA plots.

### 5.5 Differential abundance analysis with ALDEx2 using scale-reliant inference

Differentially expressed genes and pathways were identified using the ALDEx2 package in R using scale simulated random variable methods [17, 21, 23]. The underlying scale was estimated by running the ‘aldex.clr’ function with gamma set to 1e*^−^*^3^ (i.e. nearly 0). The scale values for the two conditions can be determined as the average of the rows of the ‘@scaleSamps’ slot in the resulting ‘clr‘ class object. We observed that the mean scale values differed by approximately 8-fold (∼ 2^2.95^), and that this resulted in the housekeeping functions being offset from the line of no difference (Sup-plementary Figure S1. Examining the scale of just the housekeeping functions in the same manner suggested that a scale difference of ∼ 0.15 would be more appropriate. Accordingly, all differential abundance analyses performed with the updated ALDEx2 package implemented a 15 % scale difference between groups, and a scale uncertainty of 0.5 standard deviations (as recommended previously [17, 21]). All feature tables were log_2_-transformed across 128 Monte-Carlo instances with these parameters.

All values calculated by ALDEx2 and used for analysis are the median values of the posterior distribution for each gene or function in each sample. Median scaled log-ratio (SLR) values, absolute effect sizes and expected *P* -values from Benjamini-Hochberg-corrected Welch’s t-test were calculated across all Monte-Carlo replicates. and were accessed using the ‘include.sample.summary = TRUE’ parameter within the ‘aldex.effect’ function. Appropriate effect size thresholds were assessed using the ‘aldex.plot’ function, and KO or EggNOG terms with an absolute effect size ≤-1 or ≥1 were considered differentially abundant. Note that all parts with these effect sizes also had an FDR of ≤1 %. Normalised Z-scores of effect size were calculated per-sample for pathways represented by differentially expressed KO/EggNOG terms 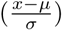 , where *x* is the median effect size, *µ* is the per-sample mean effect size and *σ* is the per-sample effect size standard deviation. Normalised Z-scores were then plotted per-pathway using the pheatmap package.

### 5.6 Identification of signature functions between groups

Signature functions were identified for groups of metatranscriptomes (healthy vs. BV and BV subgroups 1 vs. 2) using the ‘coda4microbiome’ R package. Feature tables were aggregated by KEGG pathway and subject to SLR transformation using ‘aldex.clr’, calculating the geometric mean using only the low-variance, high-abundance features.

Pathways with a median within-group dispersion ≤4 and effect score ≥0.5 were passed to the ‘coda glmnet’ function to identify signature functions.

### 5.7 Data availability and code

Raw FASTQ files are available for each of the datasets used. London and European datasets are immediately available from the European Nucleotide Archive under the accession numbers PRJEB31833 and PRJEB21446, respectively.

The Virginia dataset was obtained from the Database of Genotypes and Phenotypes (dbGaP; accession no. phs001523.v1.p1) after a successful data access request, following the conditions of the data access policy under the University of Western Ontario Research Ethics Board approval #123506.

Whole genome sequences of *Gardnerella* spp. type strains used in this study can be found under BioProject accession PRJNA474758 on NCBI. Whole genome sequences for the additional *Gardnerella* strains used in this study can be found under the following BioProject accessions: *G. leopoldii* UMB6774, PRJNA970254; *G. piotii* JNFY15, PRJNA761238; *G. swidsinskii* JNFY3, PRJNA761238; and *G. vaginalis* NR001, PRJNA394757.

All code used for the re-processing of FASTQ files at the command line and data analysis in RStudio is available as a code stable release at: https://github.com/scottdossantos/dossantos2024study.

### 5.8 Re-assigning pathways

All gene function assignments were performed automatically using the Kyoto Encyclopedia of Genes and Genomes [70]. However, due to the unique environment and somewhat generic annotation in KEGG, several assignments were clearly incorrect. Each function was individually investigated and re-assigned using the Kyoto Encyclopedia of Genes and Genomes. First the KO numbers were identified, then each function was investigated looking at the pathway, enzymatic function, chemical reaction, location, and which species was expressing it in our samples. Using that knowledge, as well as knowledge of the vaginal environment, the most probable function was inferred. KO terms that were modified, including the rationale for each, are in the ‘Supplement’ directory under ‘pathway-changes.xlsx’ and in ‘lon eur health vs BV.R’ and ‘virginia definingMolecularBV.R’ in the ‘code’ directory of the GitHub repository for this study. The latter two R scripts also contain DOIs linking to peer-reviewed literature supporting the KO term changes at the end of the files.

### 5.9 Author contributions

Author contributions are listed below, according to the CRediT taxonomy:

**Funding acquisition:** GR, GBG

**Conceptualisation:** GR, GBG

**Data curation:** SJDS, CC, JMM

**Formal analysis:** SJDS, CC, JMM, GG

**Writing-original draft:** SJDS, CC, GBG

**Writing-review & editing:** All authors

---

**Supplementary Fig S1.**
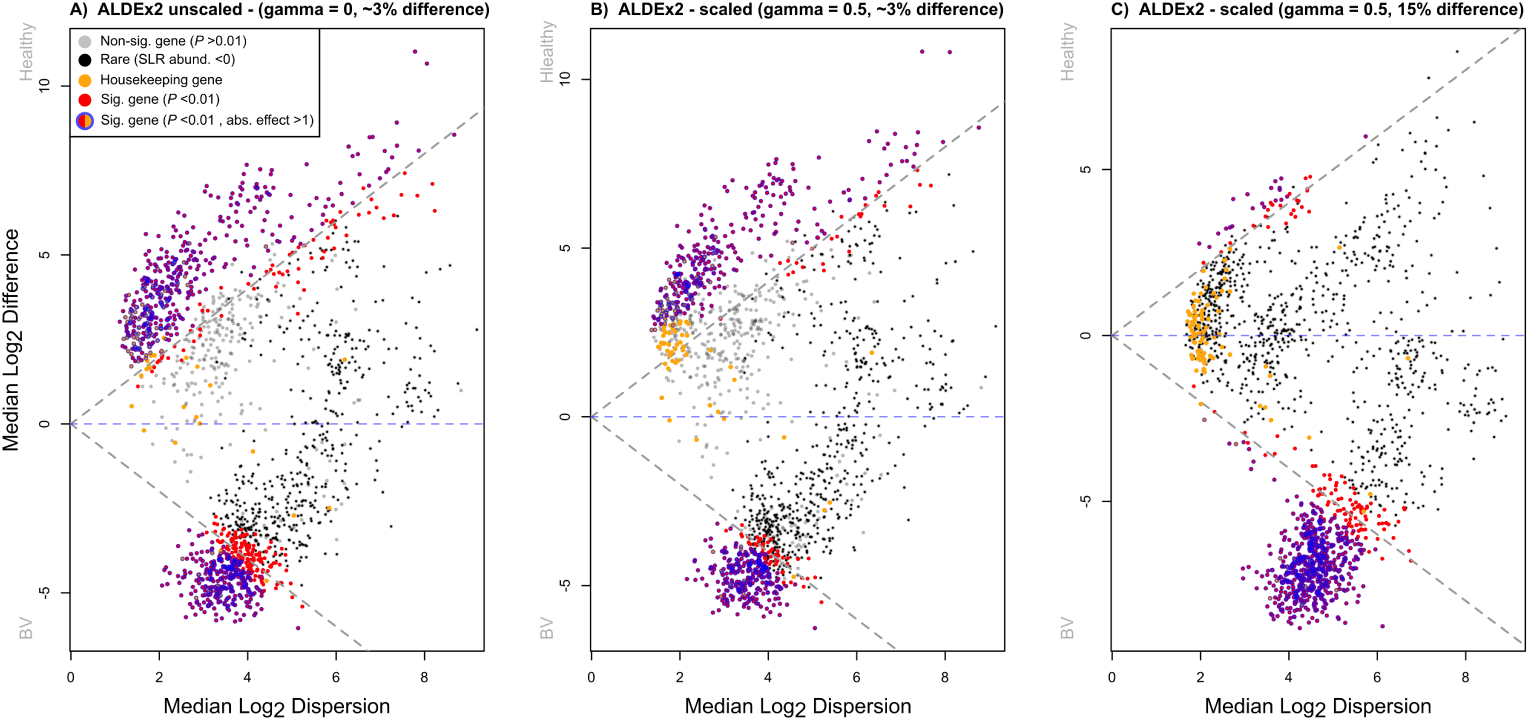
Using a between-group scale offset of 15 % is sufficient to centre a vaginal metatranscriptomic dataset: MW effect plots were produced for the London/Europe discovery dataset normalised using (**A**) ALDEx2 without scale simulation, (**B**) ALDEx2 operating with the default scale model (gamma = 0.5) or (**C**) ALDEx2 running with a full scale model (gamma = 0.5, 15 % scale offset between healthy and BV groups). Not accounting for scale results in a large number of false-positive results (**A**) and a clear error in the centre-point of the data which is, by definition, determined by the housekeeping features. Adding a small amount of scale uncertainty (**B**) increases within-group dispersion of all features such that many housekeeping genes are no longer differential; however, only the full scale model (**C**) correctly centres the housekeeping genes around the point of no difference between groups and corrects the large number of false-positive and -negative results. Colouring of data points as follows: non-significant features = grey (*P* ≥0.01), non-significant, rare features = black (median SLR abundance ¡0), housekeeping features = orange (KO terms corresponding to ’Glycolysis’, ’Ribosome’ or ’Aminoacyl-tRNA biosynthesis’), significant features based on *P*-value alone = red (*P* ≤0.01) and significant features based on *P*-value and effect size = red with blue outline (*P* ≤0.01, absolute effect size ≥1). Grey dashed lines indicate equivalence of between- and within-group difference; blue dashed line indicates no difference between groups.

**Supplementary Fig S2.**
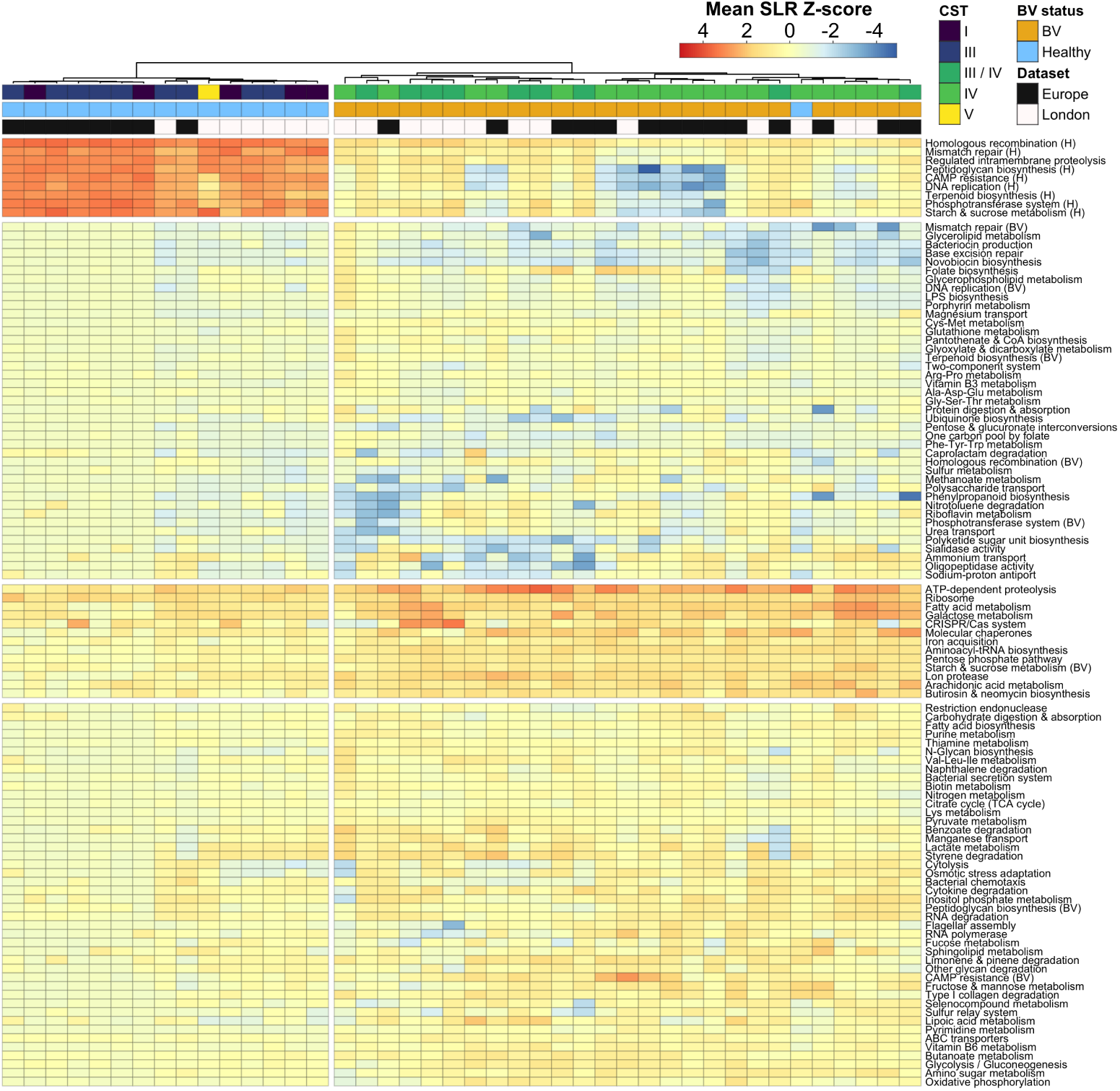
Differentially expressed pathways between health and BV – London/Europe dataset: Gene assignments were grouped by KEGG identifiers regardless of species for each sample and all differentially abundant KEGG pathways between healthy and BV metatranscriptomes in the London/Europe dataset are shown. All post-SRI absolute effect sizes and expected false discovery rates calculated by ALDEx2 are >1 and <0.01, respectively. CST, BV status and dataset indicated by colour bars.

**Supplementary Fig S3.**
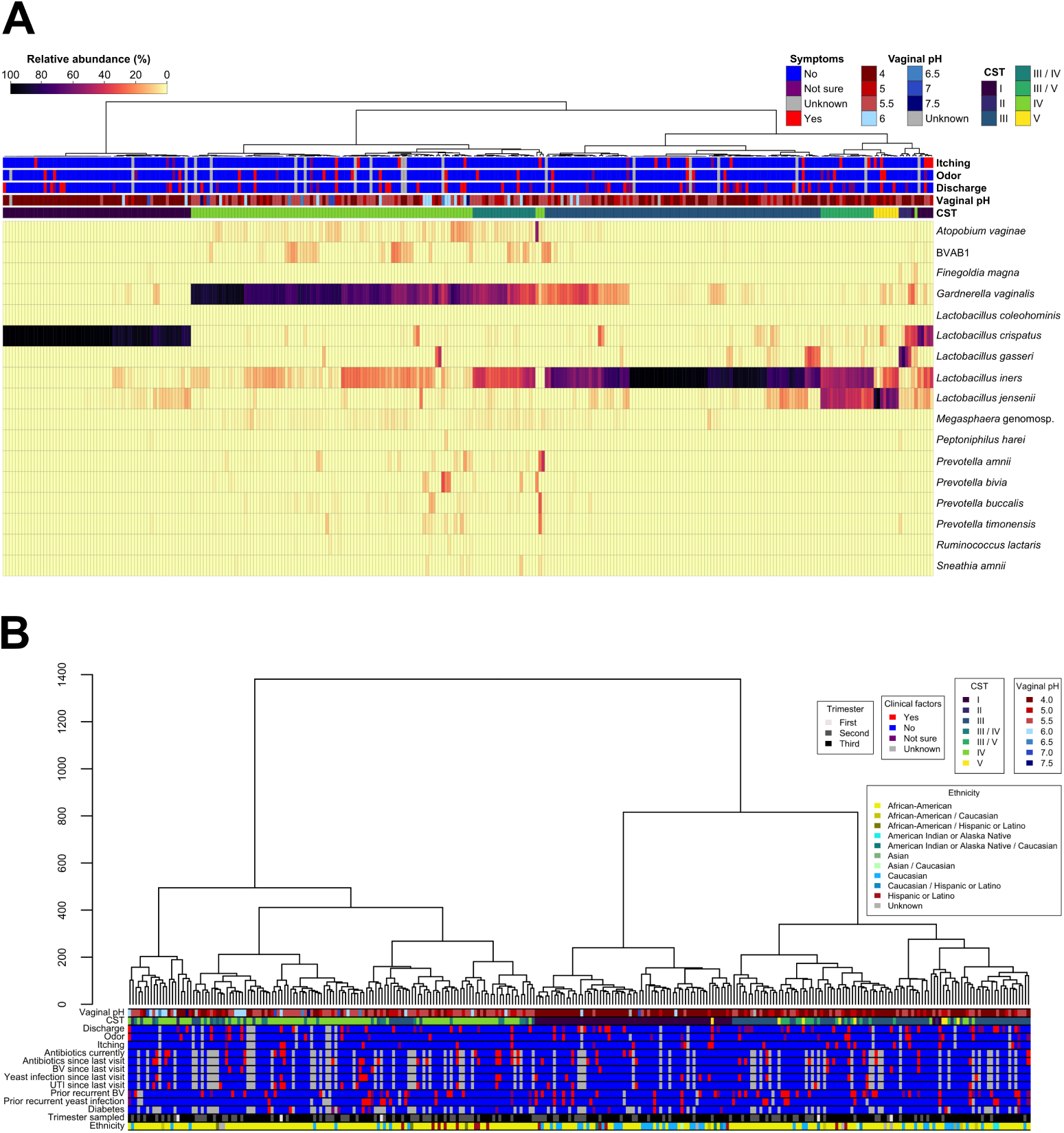
Definition of molecular states of health and BV by clustering of metatranscriptomes at the functional level: All reads from the Virginia dataset were aggregated by species (relative abundance; **A**) or function (KO term; **B**), prior to hierarchical clustering of Euclidean distances using Ward’s method. Colour bars indicate vaginal pH, CST, clinical metadata, and ethnicity. (**A**) Relative abundances of metatranscriptome reads with known taxonomy; species represented by ≥75 genes are shown. (**B**) Dendrogram of SLR-transformed, post-SRI metatranscriptome profiles aggregated by KO term, regardless of taxonomy.

**Supplementary Fig S4.**
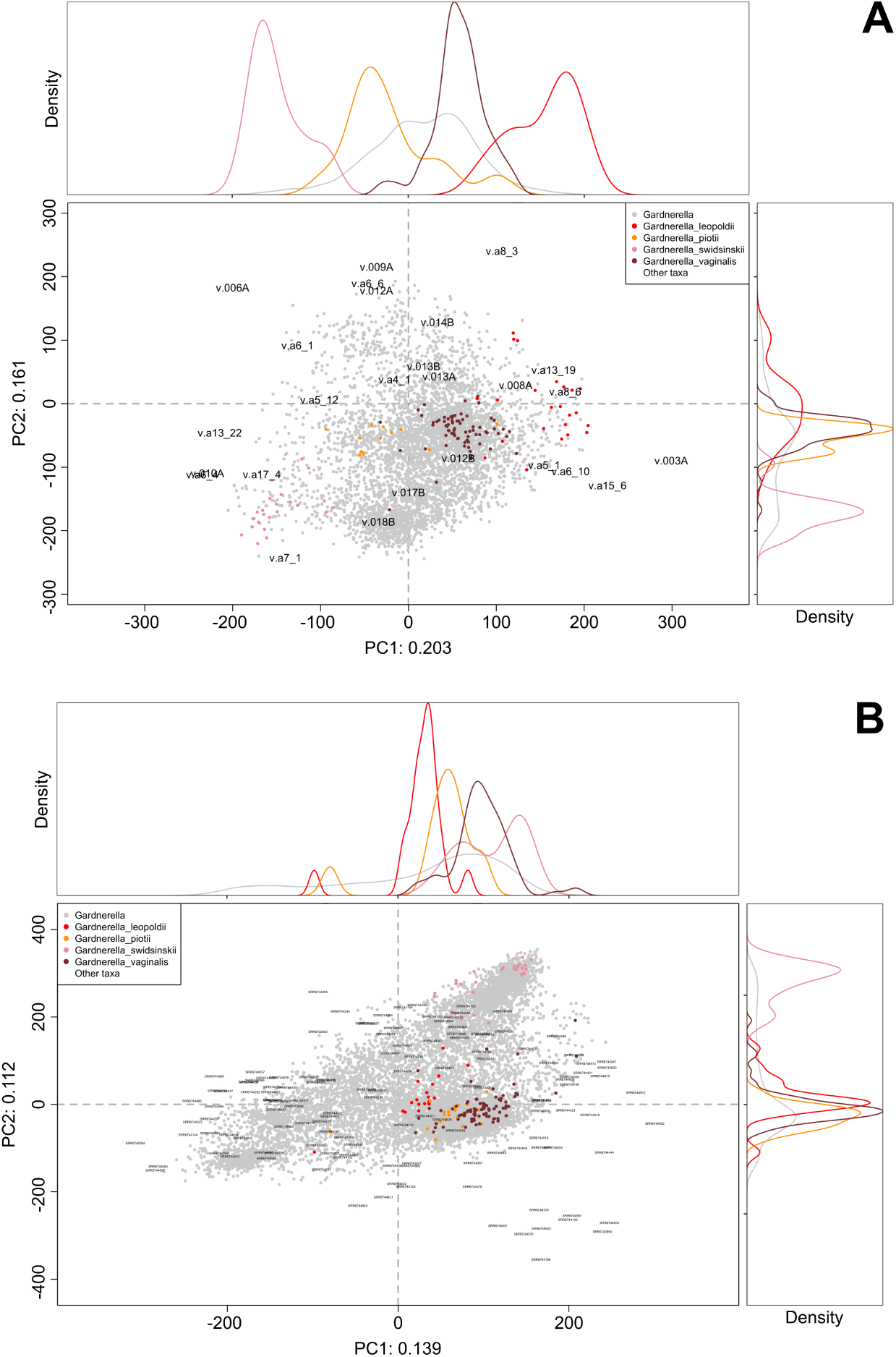
Individual *Gardnerella* species can be distinguished within meta-transcriptome profiles: Metatranscriptome profiles from London/Europe (**A**) and Virginia (**B**) datasets were subject to SLR-transformation prior to PCA. All gene features were included in the ordination; however only reads assigned to *Gardnerella* are shown. Genes found to be unique to each of the named *Gardnerella* species by pangenome analyses are coloured.

**Supplementary Fig S5.**
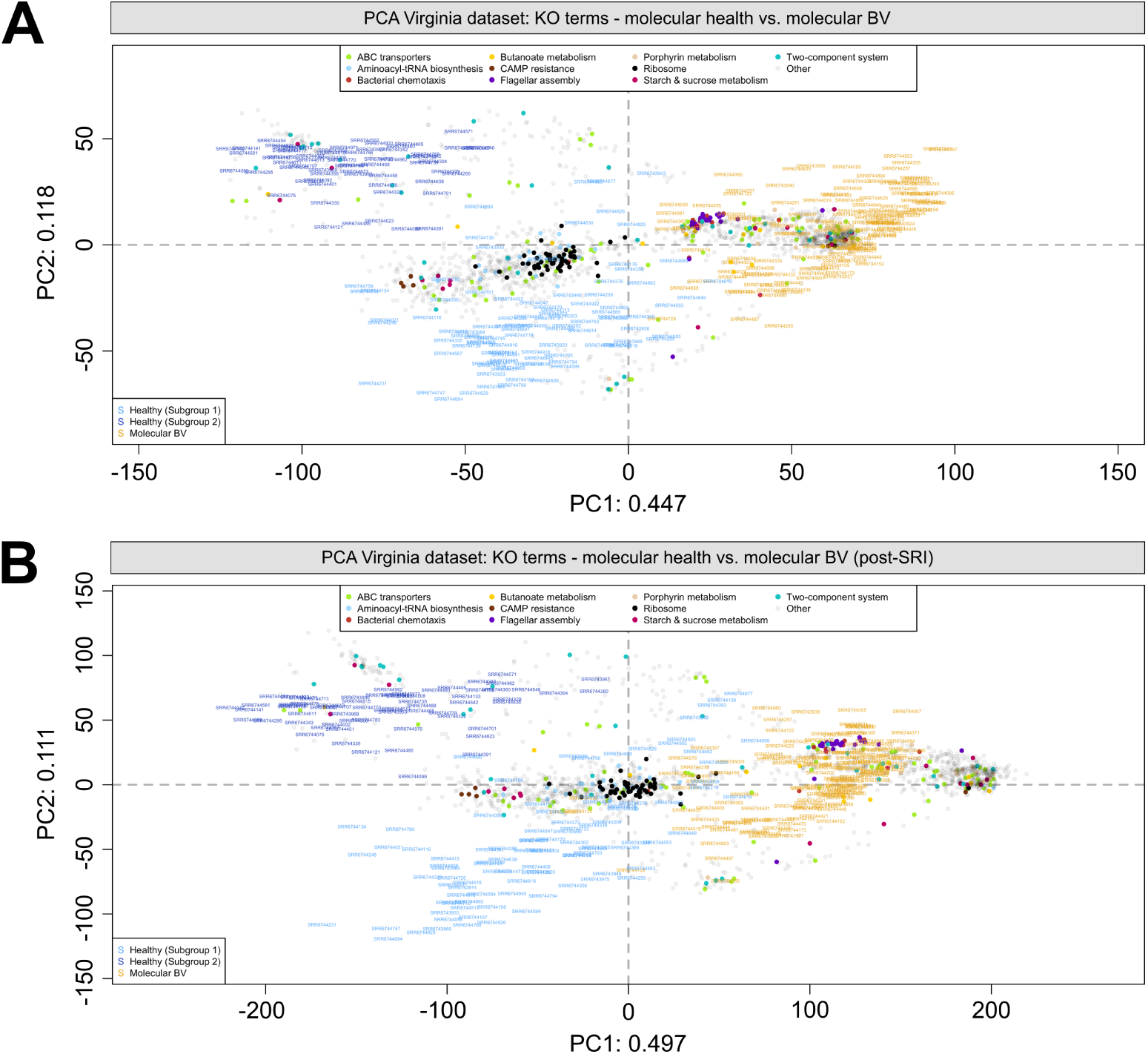
Correcting data asymmetry with SRI effectively centres genes of no-difference: KO-aggregated metatranscriptome profiles from the Virginia dataset underwent SLR-transformation either alone (**A**) or with SRI (**B**; 15 % scale difference, gamma = 0.5) prior to PCA. KO terms coloured by KEGG pathway assignments for a selection of relevant KEGG pathways. Samples coloured by group according to hierarchical clustering in Supplementary Figure S3.

**Supplementary Fig S6.**
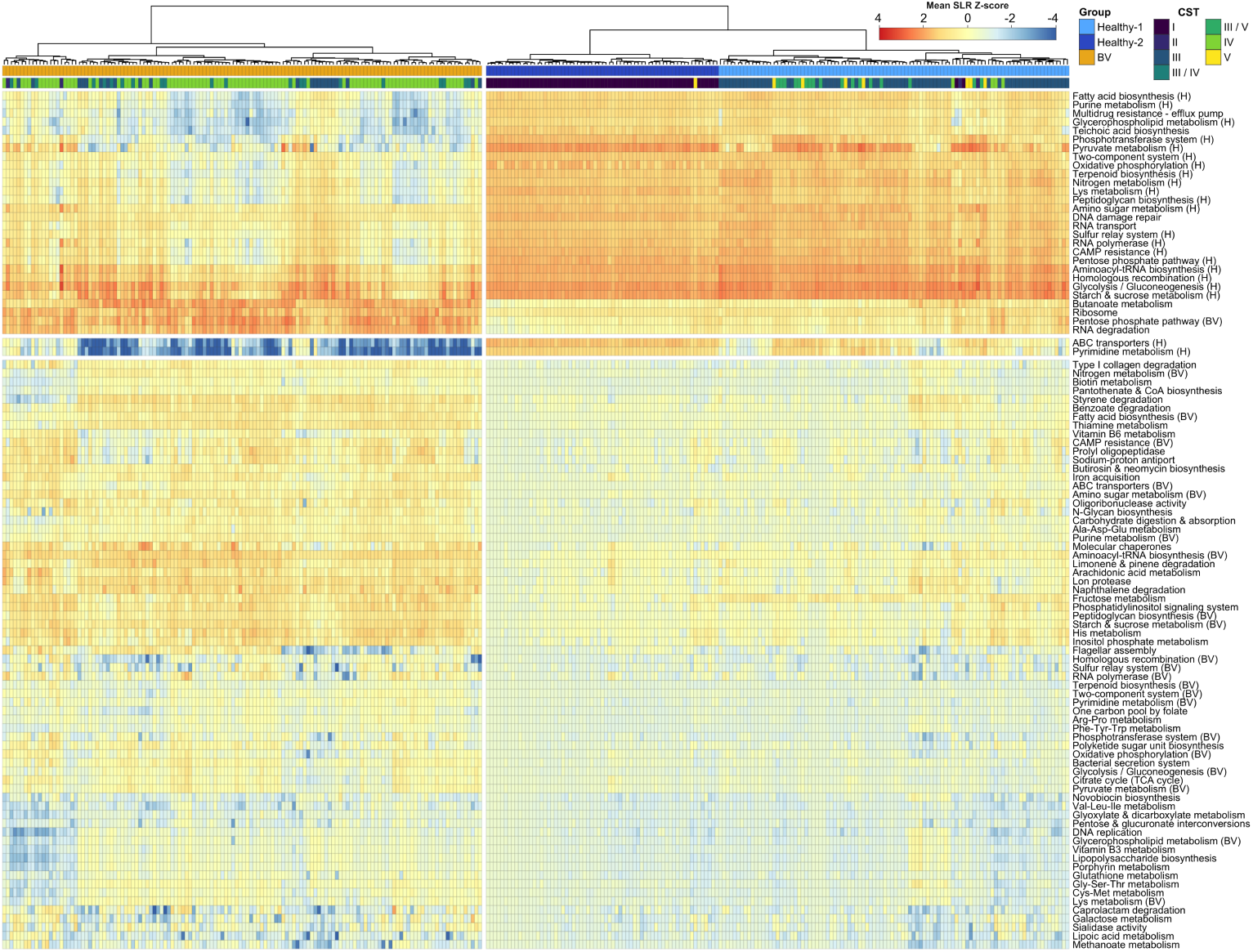
Major differentially expressed pathways of the vaginal meta-transcriptome can be replicated across datasets: Metatranscriptome profiles from the Virginia dataset were aggregated by KO term prior to SLR-transformation and SRI using ALDEx2. Analysis was stratified by states of molecular health or BV and Z-scores of mean SLR values per pathway are plotted. Colour bars indicate group and CST. All post-SRI effect sizes and BH-corrected *P*-values calculated by ALDEx2 are ≤0.01 and <1 %, respectively.

**Supplementary Fig S7.**
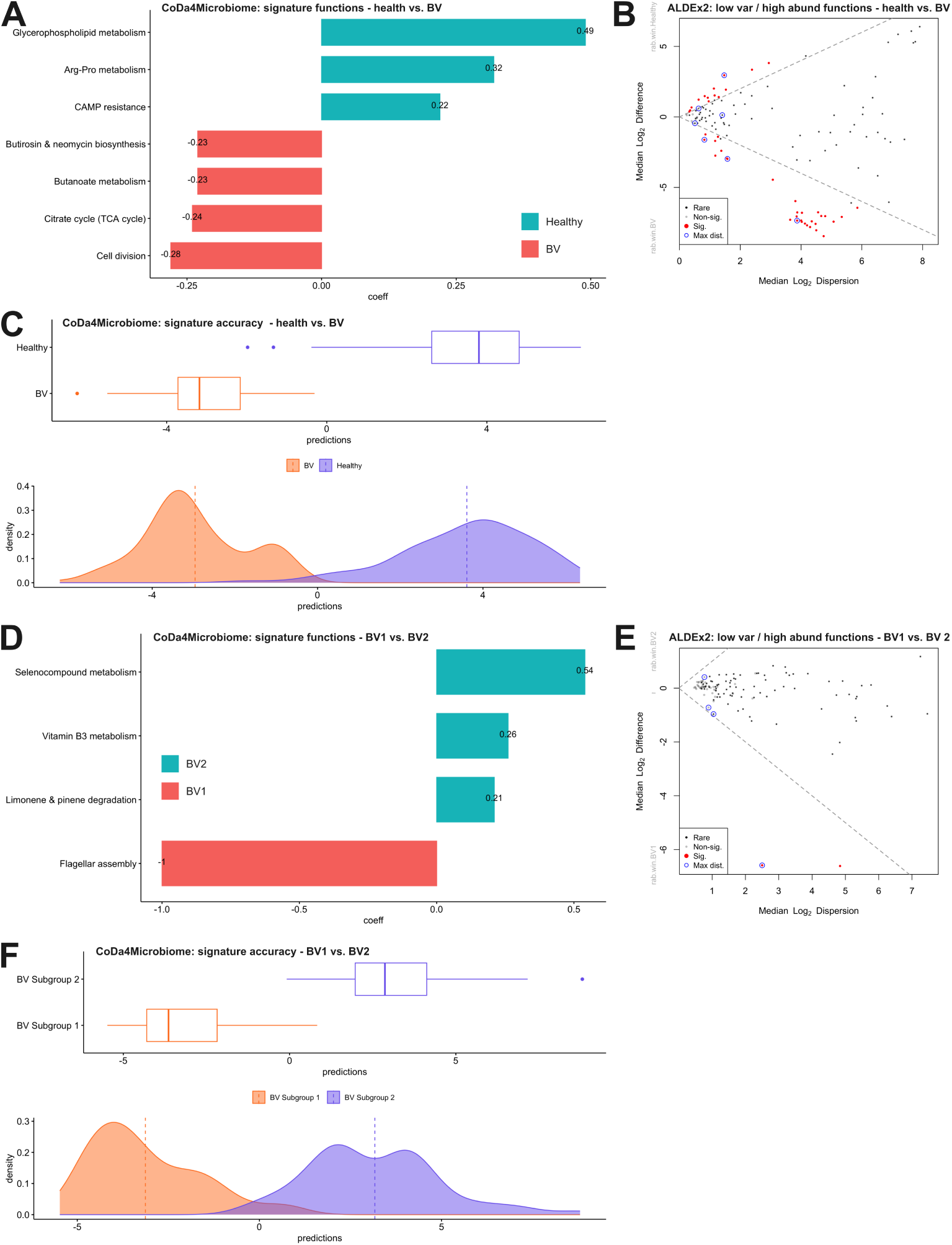
Prediction of signature functions complements differential abundance analyses: Prediction of signature functions for health vs. BV (A-C) and BV subgroups (D-F) by the ’coda4microbiome’ R package. Sets of functions identified in signature plots (A,D) exhibit high AUC values of 0.9975 and 0.9947, respectively, and are indicated on ALDEx2 effect plots in blue. Prediction plots (C,F) demonstrate a high degree of separation by group for these signatures, indicating that they define the respective groups with a high degree of accuracy. These results should be interpreted with caution, given that calculating the geometric mean from low-variance, high-abundance features does not eliminate the problematic data asymmetry shown in Suppl. Fig S1.

**Supplementary Fig S8.**
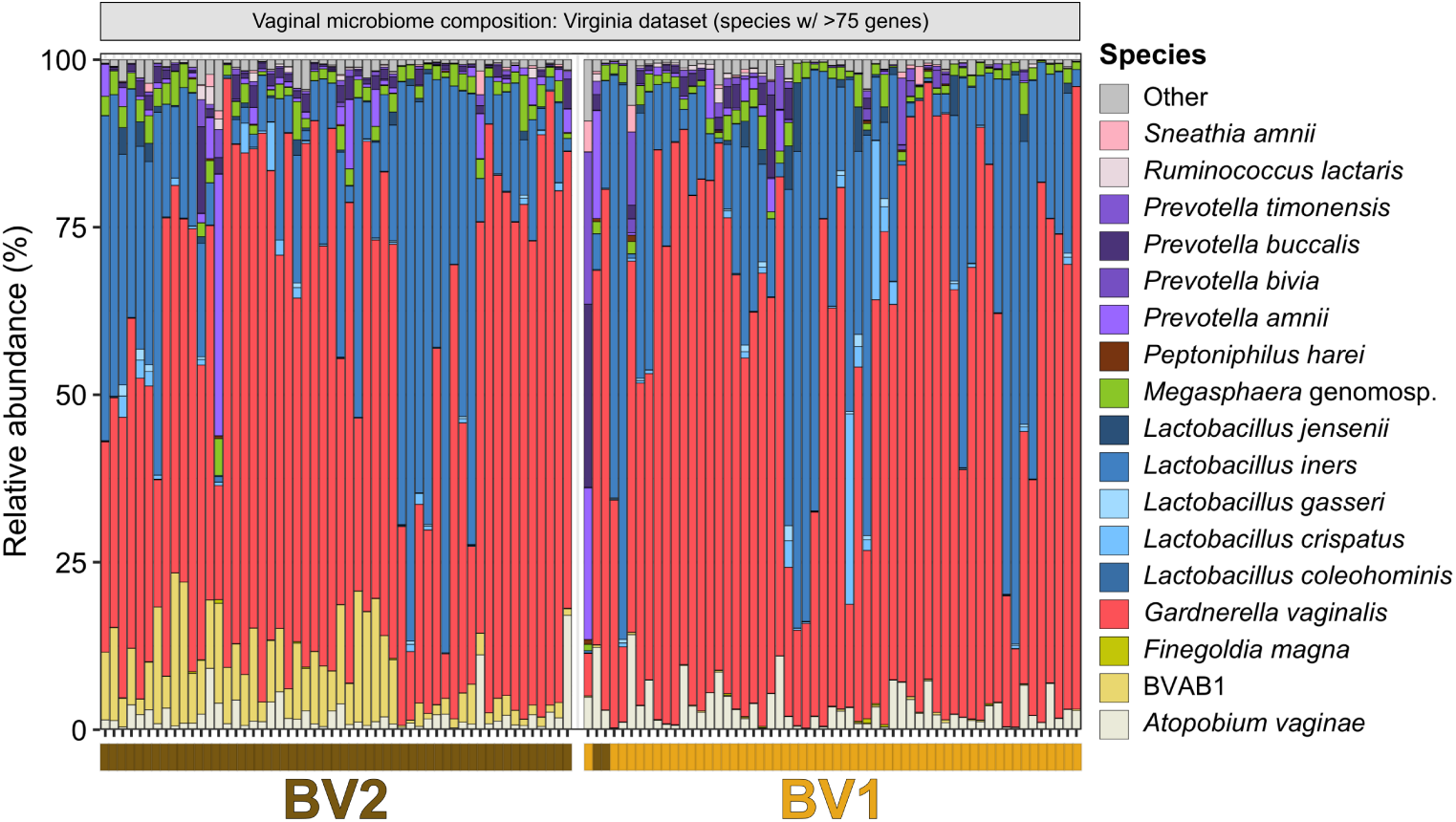
BVAB1 abundance distinguishes the two main molecular BV subgroups within the Virginia dataset: Metatranscriptome profiles of samples belonging to the two main BV subgroups within the Virginia dataset are expressed as relative abundance. Reads of known taxonomy, corresponding to species represented by ≥75 genes are included. Subgroup membership indicated below plot.

**Supplementary Fig S9.**
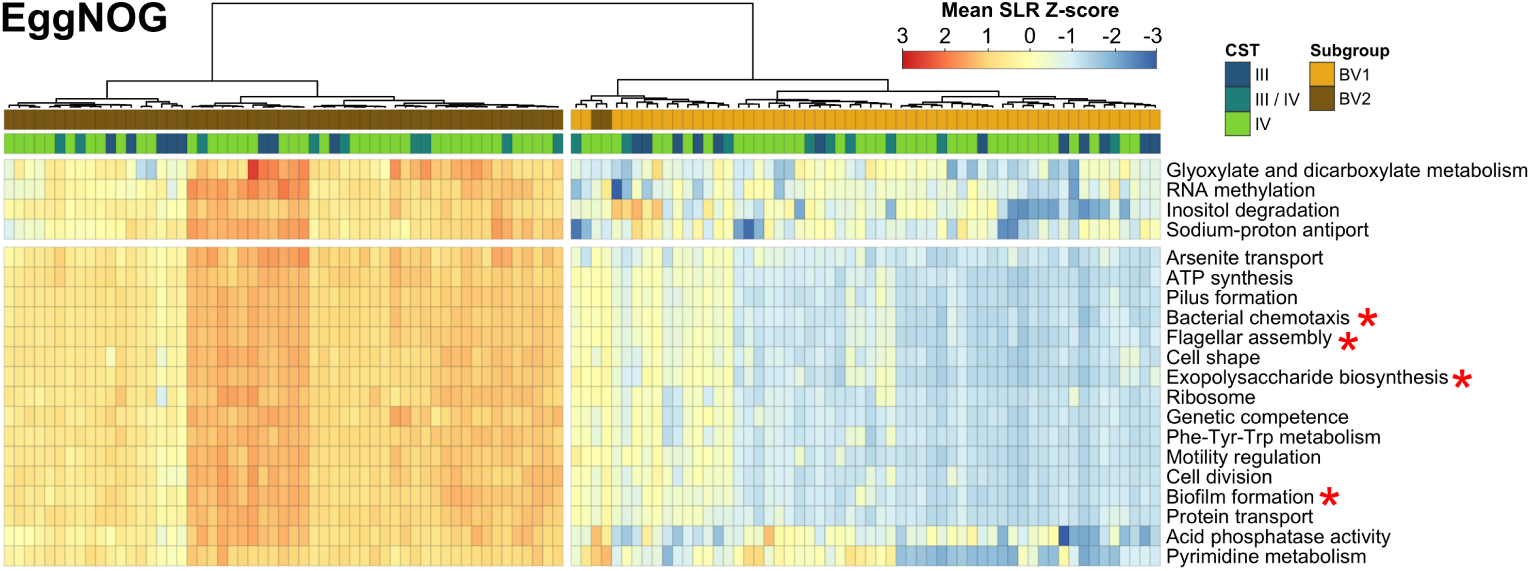
Functional differences between BV subgroups are not dependent on enzyme classification systems: Metatranscriptome profiles from the Virginia dataset were aggregated by KO term prior to SLR-transformation and SRI using ALDEx2. Analysis was stratified by membership of molecular BV subgroups and Z-scores of mean SLR values per pathway are plotted. Red asterisks indicate functions identified in the corresponding KEGG KO term analysis. Colour bars indicate group and CST. All post-SRI effect sizes and BH-corrected *P*-values calculated by ALDEx2 are ≤0.01 and <1 %, respectively.

**Supplementary Fig S10.**
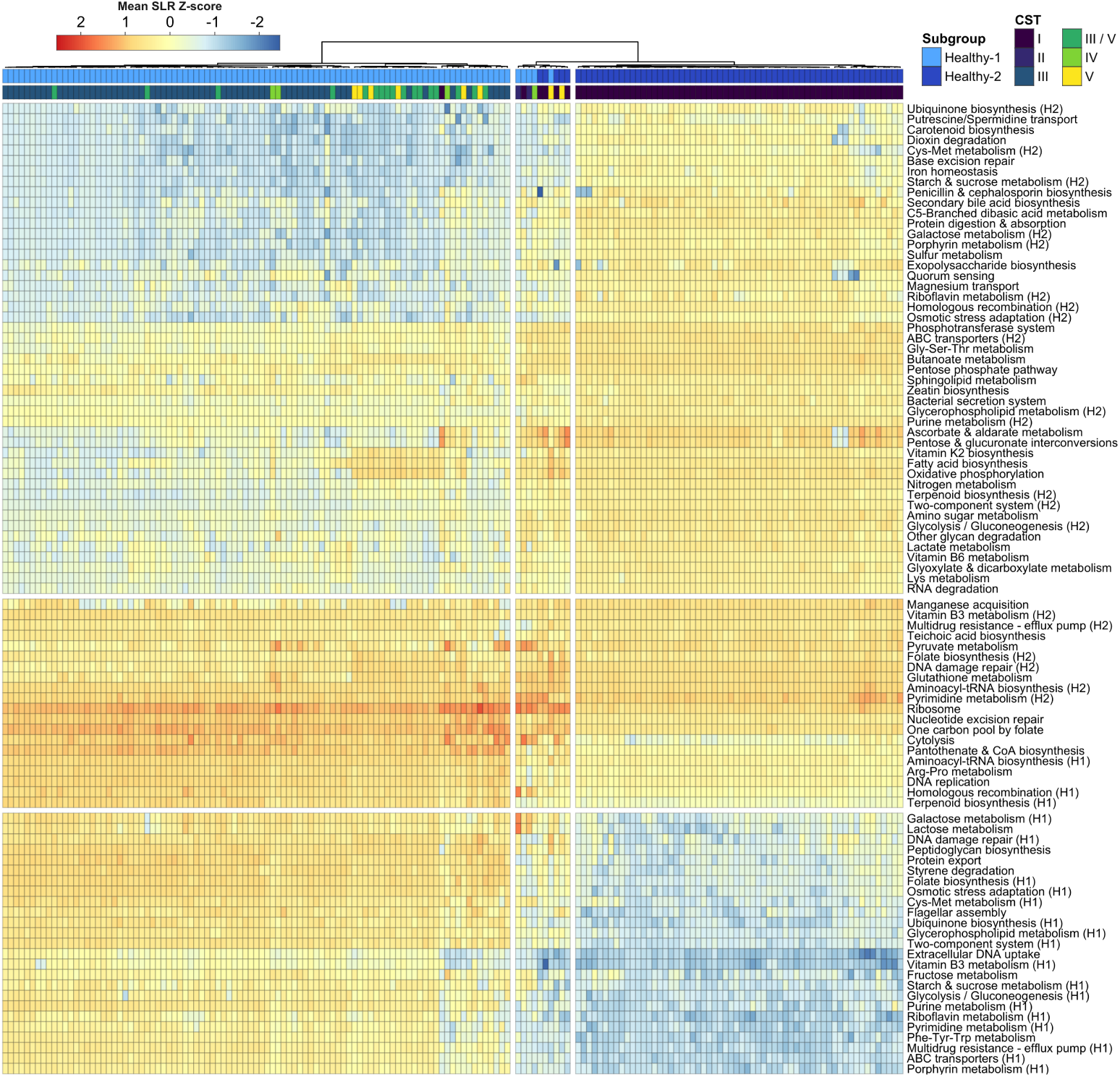
Differentially expressed pathways between molecular health subgroups –Virginia dataset: Metatranscriptome profiles from the Virginia dataset were aggregated by KO term prior to SLR-transformation and SRI using ALDEx2. Analysis was stratified by membership of molecular health subgroups and Z-scores of mean SLR values per pathway are plotted. Colour bars indicate group and CST. All post-SRI effect sizes and BH-corrected *P*-values calculated by ALDEx2 are ≤0.01 and <1 %, respectively.

## Notes

https://github.com/scottdossantos/dossantos2024study

### Competing Interest Statement

The authors have declared no competing interest.

### Summary of Updates

Removed correlation analysis of gestational age at delivery and vaginal metatranscriptome data at the taxonomic/functional levels. We are not able to share this data (which originates from the MOMS-PI study) in a public repository, which violates the data access policy of the journal the manuscript has been submitted to. Removing this analysis does not alter any of the conclusions we can draw from our study; therefore, we have removed this analysis from the article to comply with the journal's standards on reproducibility and data access.

